# Mouse and cellular models of *KPTN*-related disorder implicate mTOR signalling in cognitive and progressive overgrowth phenotypes

**DOI:** 10.1101/2022.07.15.500213

**Authors:** Maria O. Levitin, Lettie E Rawlins, Gabriela Sanchez-Andrade, Osama A. Arshad, Stephan C. Collins, Stephen J. Sawiak, Phillip H. Iffland, Malin H.L. Andersson, Caleb Bupp, Emma L. Cambridge, Eve L. Coomber, Ian Ellis, Johanna C. Herkert, Holly Ironfield, Logan Jory, Perrine F. Kretz, Sarina G. Kant, Alexandra Neaverson, Esther Nibbeling, Christine Rowley, Emily Relton, Mark Sanderson, Ethan M. Scott, Helen Stewart, Andrew Y. Shuen, John Schreiber, Liz Tuck, James Tonks, Thorkild Terkelsen, Conny van Ravenswaaij-Arts, Pradeep Vasudevan, Olivia Wenger, Michael Wright, Andrew Day, Adam Hunter, Minal Patel, Christopher J. Lelliott, Peter B. Crino, Binnaz Yalcin, Andrew Crosby, Emma L. Baple, Darren W. Logan, Matthew E. Hurles, Sebastian S. Gerety

**Affiliations:** Wellcome Sanger Institute, Wellcome Genome Campus, Hinxton, Cambridge, UK; Evox Therapeutics Limited, Oxford, UK; College of Medicine and Health, University of Exeter, Exeter, UK; INSERM Unit 1231, Université de Bourgogne Franche-Comté, Dijon, France; Behavioural and Clinical Neuroscience Institute, University of Cambridge, Cambridge, UK; Wolfson Brain Imaging Centre, Department of Clinical Neurosciences, University of Cambridge, Cambridge, UK; Dept of Neurology University of Maryland School of Medicine, Baltimore, MD, USA; Spectrum Health, Helen DeVos Children’s Hospital, Grand Rapids, MI, USA; Liverpool Women’s NHS Foundation Trust, Alder Hey Children’s Hospital, Liverpool, UK; Department of Genetics, University Medical Centre, University of Groningen, Groningen, the Netherlands; Haven Clinical Psychology Practice Ltd, Bude, Cornwall, UK; IGBMC, UMR7104, Inserm, Illkirch, France; Department of Clinical Genetics, Erasmus MC, University Medical Center Rotterdam, Rotterdam, the Netherlands; Department of Clinical Genetics, Leiden University Medical Center, Leiden, the Netherlands; Open Targets, Wellcome Genome Campus, Hinxton, Cambridge, UK; Department of Genetics, University of Cambridge, Cambridge, UK; Laboratory for Diagnostic Genome Analysis, Department of Clinical Genetics, Leiden University Medical Center, Leiden, the Netherlands; Institute of Metabolic Science, Cambridge University, Cambridge, UK; Faculty of Health and medical science, University of Surrey, Guildford, UK; New Leaf Center, Clinic for Special Children, Mount Eaton, OH, USA; Oxford Centre for Genomic Medicine, Oxford University Hospitals NHS Trust, Oxford, UK; London Health Sciences Centre, London, ON, Canada.; Division of Medical Genetics, Department of Pediatrics, Schulich School of Medicine and Dentistry, Western University, London, ON, Canada; Department of Neurology, Children’s National Medical Center, Washington DC, USA; Department of Clinical Genetics, Aarhus University Hospital, Aarhus, Denmark; Department of Clinical Genetics, University Hospitals of Leicester, Leicester Royal Infirmary, Leicester, UK; Institute of Human Genetics, International Centre for Life, Newcastle upon Tyne, UK; Qkine Ltd., Cambridge, UK; Waltham Petcare Science Institute, Waltham on the Wolds, UK

## Abstract

*KPTN*-related disorder (KRD) is an autosomal recessive disorder associated with germline variants in *KPTN* (*kaptin*), a component of the mTOR regulatory complex KICSTOR. To gain further insights into the pathogenesis of KRD, we analysed mouse knockout and human stem cell KPTN loss-of-function models. *Kptn*^−/−^ mice display many of the key KRD phenotypes, including brain overgrowth, behavioural abnormalities, and cognitive deficits. Assessment of affected individuals has identified concordant selectivity of cognitive deficits, postnatal onset of brain overgrowth, and a previously unrecognised KPTN dosage-sensitivity, resulting in increased head circumference in their heterozygous parents. Molecular and structural analysis of *Kptn*^−/−^ mice revealed pathological changes, including differences in brain size, shape, and cell numbers primarily due to abnormal postnatal brain development. Both the mouse and differentiated iPSC models of the disorder display transcriptional and biochemical evidence for altered mTOR pathway signalling, supporting the role of KPTN in regulating mTORC1. Increased mTOR signalling downstream of KPTN is rapamycin sensitive, highlighting possible therapeutic avenues with currently available mTOR inhibitors. These findings place KRD in the broader group of mTORC1 related disorders affecting brain structure, cognitive function, and network integrity.

## Introduction

Human brain development requires an intricate and tightly regulated sequence of events involving cell proliferation, migration, differentiation, and integration into cohesive circuitry^1^, complex processes that are highly sensitive to changes in gene expression. Neurodevelopmental disorders (NDDs) with underlying genetic causes have been associated with most stages of brain development^2^. Prevalent phenotypic manifestations include changes in brain volume and cranial abnormalities^3–5^. A minority of these size changes involve head overgrowth (e.g. 1250 out of 5255, or 24%, of cases in the Decipher open access patient collection^6^ where a head size change was observed), termed *macrocephaly* where the occipitofrontal circumference (OFC) exceeds 2 standard deviations (SD). Megalencephaly is typically used to describe such overgrowth when it specifically affects brain structures (neurons, glia, ventricles), rather than simply the anatomical dimensions of the head and skull^7,8^.

For overgrowth disorders, there is a frequent convergence on a smaller set of growth regulating pathways, including AKT and mTOR (mechanistic target of rapamycin) signalling^9^. mTOR, a serine/threonine kinase that forms the catalytic subunit of two distinct protein complexes known as mTOR Complex 1 (mTORC1) and 2 (mTORC2)^10–13^, regulates protein synthesis, cell growth, and nutrient uptake. Moreover, mTOR signalling is central to the regulation of long-lasting synaptic plasticity, which in turn is critical to the formation and persistence of memories^14^. Aberrant mTOR signalling is associated with numerous diseases such as cancer, diabetes, and neurological disorders including epilepsy, autism, developmental delay and macrocephaly^10–13^. *De novo* and inherited variants have been identified in numerous mTOR pathway genes, leading to the term “mTORopathies” to describe the involvement of the mTORC1 signalling pathway in these disorders^15–20^. A growing body of literature is uncovering the intricate relationship between the mTOR pathway, neural stem cell proliferation, and neuronal differentiation^21–24^, providing insights into the mechanisms of malformations of cortical development such as tuberous sclerosis complex, focal cortical dysplasia, hemimegalencephaly, and megalencephaly^25–27^.

We previously described inherited biallelic variants in the *KPTN* gene as underlying KRD, also known as **M**acrocephaly, **A**utistic features, **S**eizures, **D**evelopmental delay (MASD) syndrome, (OMIM 615637), an autosomal recessive neurodevelopmental disorder in nine affected individuals from an extended interrelated family of Ohio Amish background^28^. All affected individuals were either homozygous or compound heterozygous for two *KPTN* founder variants ((p.(Ser259*) and p.(Met241_Gln246dup)). The cardinal clinical features of KRD include craniofacial dysmorphism (frontal bossing, long face with prominent chin, broad nasal tip and hooded eyelids), macrocephaly with OFC measurements up to 5.4 SD above the mean, hypotonia in infancy, global developmental delay, intellectual disability (mild to severe), and behavioural features including anxiety, hyperactivity, stereotypies and repetitive speech. Other more variable features included seizures (in 1/3, onset between 3 months and 7 years), splenomegaly and recurrent respiratory tract infections. Where available, neuroimaging revealed a globally enlarged but structurally normal brain. Subsequent studies have identified six additional affected non-Amish individuals with biallelic *KPTN* variants and a similar clinical presentation. To date five distinct variants in *KPTN* have been published^29–32^ that were identified in KRD probands.

KPTN has been identified as a negative upstream regulator of the mTORC1 signalling pathway as part of the KICSTOR protein complex that also includes SZT2, ITFG2, and C12orf66. In cell lines, KICSTOR is required for the inhibition of mTORC1 signalling in response to amino acid or glucose deprivation^11,33^. KPTN forms a heterodimer with ITFG2, which then complexes with SZT2. Interestingly, biallelic variants in *SZT2* also cause a distinct NDD, characterised by severe early-onset epileptic encephalopathy, global developmental delay, structural brain abnormalities, and frequent macrocephaly^17,18^. In mice, *Szt2* LoF mutants display seizure susceptibility and increased mTORC1 signalling in the brain^15,34^.

We aimed to assess the functional consequences of homozygous *Kptn* LoF in mice (referred to herein as *Kptn* KO mice), as well as human iPSC models of the condition, as to date no study has modelled KRD *in vivo* or *in vitro*. Our mouse model recapitulates the primary human phenotypes, including cognitive and behavioural impairments and skull and brain overgrowth. Molecular findings include the misexpression of numerous seizure-associated genes, progenitor markers, as well as biochemical and transcriptional evidence for increased mTOR signalling. The aberrant mTOR signalling is corrected by acute treatment with rapamycin, a known mTOR inhibitor, supporting a pre-clinical path to therapeutic intervention. Cortical neural precursors differentiated from human iPSCs null for *KPTN* robustly support the mouse findings. We also identify a previously unrecognised effect of head overgrowth in heterozygous carriers of KPTN alleles, consistent with the recent studies showing that heterozygosity of alleles for other recessive disorders may be contributing to incompletely penetrant phenotypes in the general population^35–38^. Together, these models confirm *KPTN* to be among a growing list of mTOR modulating disease associated genes and provide a platform for further exploring the pathomechanistic basis of KRD.

## Results

### Generation and characterisation of a KPTN mouse model

A number of variants associated with KRD introduce a frame-shift resulting in premature termination and likely inactivate KPTN protein function^31^. We therefore used an engineered LoF mouse model^39^. The allele introduces a splice-trapping cassette into an intron downstream of coding exon 8, resulting in an in-frame LACZ open reading frame and premature termination of transcription via an exogenous polyadenylation signal. The resulting transcript closely resembles the truncating allele (p.(Ser259*)) observed in the original KRD publication^28^ (Supplementary Fig. 1a) which also led to premature termination in exon 8. Homozygous *Kptn* KO mice are observed at expected near-Mendelian ratios and appear to have normal body shape. Read count quantification in bulk RNA-Seq shows a robust reduction in *Kptn* mRNA pre- and postnatally, and across adult brain regions (87-90% decrease, Supplementary Fig. 1b). This confirms that the mouse model is functioning efficiently as a LoF allele, and leads to nonsense mediated decay of *Kptn* mRNA *in vivo*.

### *Kptn^−/−^* mice have increased locomotor activity and anxiety-like phenotypes

The majority of individuals with KRD display consistent neurobehavioural traits, including intellectual impairment and features suggestive of autistic spectrum disorder (anxiety, stereotypies and repetitive speech) with variable hyperactivity and seizures^28–32^. To investigate whether our mouse model recapitulates any of the neurobehavioural features of the human disorder, we evaluated *Kptn* KO mice with a series of functional tests. Locomotor capabilities, including distance covered and time spent moving, were assessed in *Kptn^−/−^* mice compared to wild-type controls, using video-assisted observation and scoring in an open-field test^40^. *Kptn^−/−^* mice travelled a significantly greater distance than *Kptn^+/+^* controls, accounted for by the increase in time spent moving (p<0.05, Fig. 1a,b). All other activity parameters were not significantly different between genotypes (data not shown). Anxiety-related behaviours are a phenotype observed in over 50% of individuals with KRD^31^. To assess anxiety in *Kptn^−/−^* mice, we performed a light/dark box test^41,42^. *Kptn^−/−^* mice spent significantly more time in the dark zone (59.6% increase, p=4.8×10^−5^), and had reduced frequency of visits to the light zone (45.6% reduction, p=0.0006, Fig. 1c,d respectively). These results indicate hyperactivity and a strong anxiety-like phenotype in the *Kptn^−/−^* mouse model, concordant with those observed in KRD^28–32^.

**FIGURE 1.**
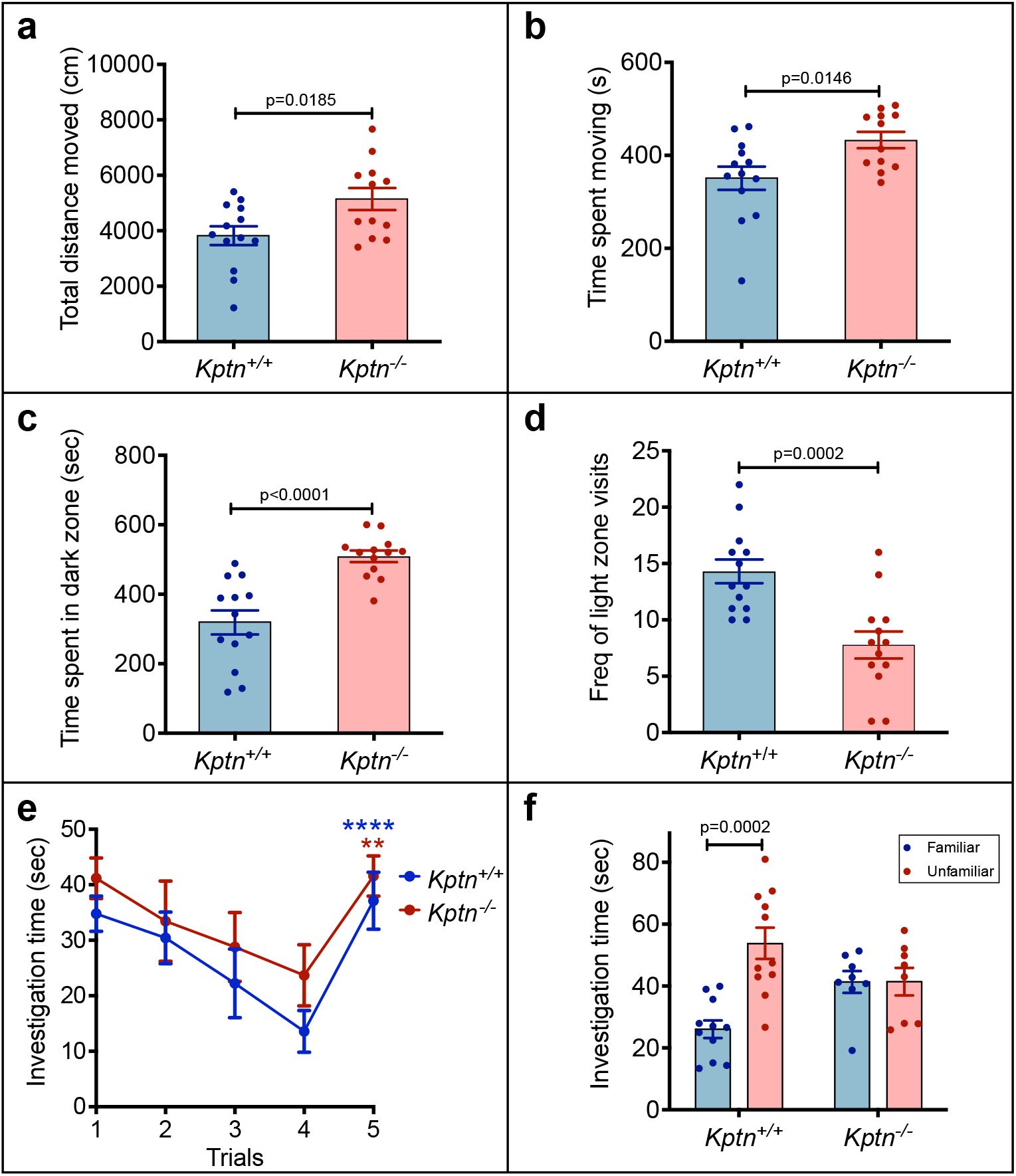
*Kptn*^−/−^mice display increased activity, anxiety and memory deficits. *Kptn*^−/−^ mice display increased locomotor activity (**a,b**) anxiety-like behaviour (**c,d**), and reduced memory retention (**e,f**). **a,b** The distance covered in an open field and time spent moving. **a** *Kptn*^−/−^ mice (n=12) covered significantly more distance (p=0.0185, t=2.534 df=23, two-tailed Student’s t-test) than wild-type controls (*Kptn*^+/+^, n=13) and **b** spent more time moving (p=0.0146, t=2.64 df=23, two-tailed Student’s t-test). **c** *Kptn*^−/−^ mice (n=13) spend significantly longer time in the dark zone (p< 0.0001; t=4.946 df=24, two-tailed Student’s t-test) of a light/dark box than wild-type controls (*Kptn* ^+/+^, n=13), and **d** have reduced frequency of visits to the light zone (p=0.0002, t=4.326 df=25, two-tailed Student’s t-test). **e** Social recognition (Day 1). Both controls (*Kptn* ^+/+^, n=11) and *Kptn* mutant mice (*Kptn^−/^*^−^, n=8) recognize a stimulus animal repeatedly presented to them over the course of four trials, as shown by a decline in the investigation time over trials 1-4. Both mutant and wild-type mice display an increase in the investigation time on trial 5 when presented with a novel stimulus animal (two-way ANOVA, interaction TRIAL x GENOTYPE F (4, 68) = 0.3852, P=0.8185; TRIAL F (4, 68) = 16.26, P<0.0001; post-hoc *Kptn*^+/+^ p<0.0001****, *Kptn*^−/−^ p=0.0023**). **f** Social discrimination (Day 2). Wild-type *Kptn*^+/+^ controls (n=11), but not *Kptn*^−/−^ mice (n=8) show a significant increase in time spent investigating an unfamiliar stimulus versus the familiar stimulus mouse from Day 1, suggesting that *Kptn^−/^*^−^ mice do not retain the memory of a familiar animal at 24 h (*Kptn*^+/+^: p=0.0002, t= 4.79 df=2; *Kptn^−/^*^−^: p=0.985, t=0.0185 df=14, two-tailed multiple t-test with multiple comparison correction). Values are plotted as mean ± SEM.

### *Kptn*^−/−^ mice have cognitive deficits

KRD in humans is characterised by highly penetrant intellectual disability. Thus, we evaluated whether *Kptn^−/−^*mice display concordant intellectual deficits. First, an ethologically relevant social recognition (SOR) assay was used to assesses olfactory-mediated hippocampus-dependent memory^43–45^. On day 1, *Kptn^−/−^* mice display no detectable deficit in social interaction, as measured by the time spent investigating the novel conspecific (stimulus A) on trial 1 (Fig. 1e). With repeated exposure over four trials, both wild-type and *Kptn^−/−^*mice habituated to the stimulus animal, indicating a functional olfactory system. Upon presentation of a novel stimulus animal, both genotypes significantly increased their investigation time (trial 5), indicating successful recognition of the novel mouse (Fig. 1e). On day 2, when given the choice between investigating an unfamiliar mouse and a familiar one (stimulus A from day 1), wild-type mice spent significantly more time investigating the unfamiliar mouse, as expected (p=0.0002, Fig. 1f). *Kptn^−/−^* mice however, did not show a preference for either stimulus (n.s., Fig. 1f), and had a significantly reduced preference index for the unfamiliar stimulus (53.9%, p=0.0136) when compared to that of *Kptn^+/+^* mice (67.5%). The results of the SOR assay indicate a memory deficit in *Kptn^−/−^*mice.

To further characterise the memory deficit, *Kptn* −/− mice were tested in a spatial memory assay, the Barnes maze^46–48^, which is similar in principle to the Morris water maze and tests a mouse’s ability to identify an escape box using spatial cues in an open arena. During both training periods (Training 1 and 2), there was no difference between *Kptn^−/−^* and wild-type mice in the time taken to approach the escape box (primary latency to approach), indicating that both genotypes were able to locate the target zone with equivalent speed (Fig. 2a). During the first probe test, after 24 h (when the escape box was removed), the time spent around the target hole relative to other holes was measured (Fig. 2b). Both genotypes spent longer around the target hole quadrant when compared to other quadrants (p≤0.05). However, during the second probe test (72 h after the end of Training 2), the *Kptn^−/−^* mice do show a deficit in long-term memory, and were on average further from the goal box location (Fig. 2c, 12.8% further, p=0.0014), and spent significantly less time in the target zone than wild-type controls (Fig. 2d, 17.9% less time, p=0.0150).

**FIGURE 2.**
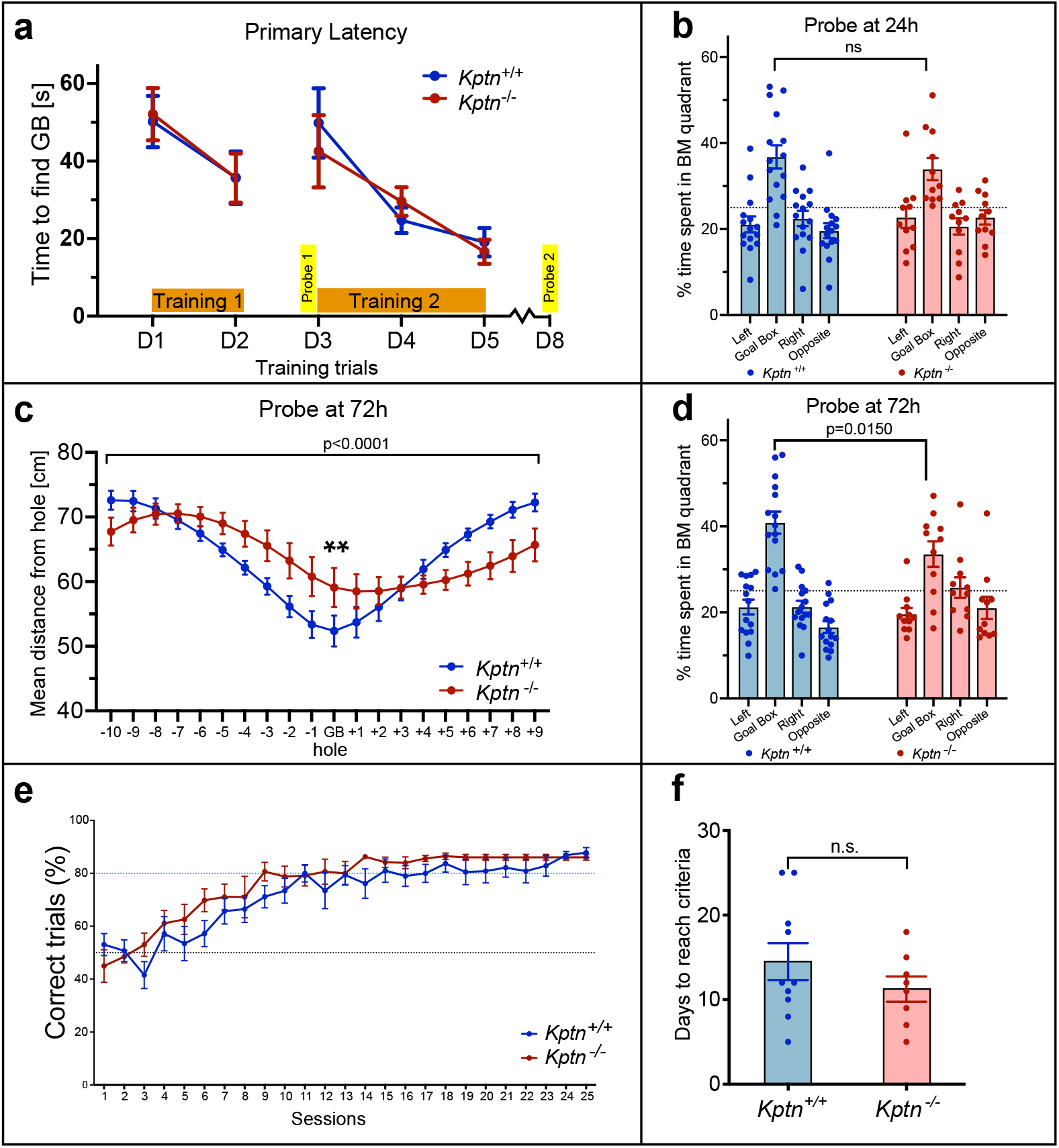
*Kptn*^−/−^ mice have specific memory deficits. **a-d** Barnes maze reveals impaired spatial memory acquisition in *Kptn^−/−^* mice**. a** Primary latency across all days of training (Training 1, D1-D2; Training 2, D3-D5) for *Kptn*^+/+^ (n=15) and *Kptn^−/−^* (n=11) mice. There was no genotype difference across all training days and a reduction in time taken to get to the escape box in Training 1 and Training 2 (2-way ANOVA, interaction GENOTYPE X ZONE F(4,120)=0.2577, p=0.9044). **b** The percentage of time *Kptn*^+/+^ (blue, n=15) and *Kptn^−/−^* (red, n=11) mice spent around the target hole quadrant (Goal Box) and non-target quadrants of the Barnes maze during a probe trial 24 h after Training 1 was similar (2-way ANOVA, interaction GENOTYPE X ZONE F(3,96)=0.8544, p=0.4676). Both genotypes spent significantly more time near the target versus all other holes (p≤0.05). **c** The mean distance from each hole during probe trial, 72 h after Training 2. *Kptn^−/−^* mice were on average further from the target hole compared to *Kptn*^+/+^ mice (2-way ANOVA, interaction GENOTYPE X HOLE F(19,500)=5.351, p=<0.0001; post-hoc FDR q=0.0045, p=0.0014**). **d** The percentage of time *Kptn*^+/+^ (blue, n=15) and *Kptn^−/−^* (red, n=11) mice spent in the target hole quadrant (Goal Box) and non-target quadrants of the Barnes maze during the probe trial, 72 h after training 2. The *Kptn^−/−^* mice spent significantly less time in the target hole quadrant than the *Kptn*^+/+^ mice (2-way ANOVA, interaction GENOTYPE X ZONE F(3,96)=3.667, p=0.0150; post-hoc FDR q=0.0632, p=0.0150) **e,f** *Kptn^−/−^* mice do not display an impaired performance in a hippocampus-independent pairwise discrimination task. **e** Percentage of correct trials (when CS+ image was nose-poked) out of the total trials completed per session for *Kptn^−/−^* (n=8) and *Kptn*^+/+^ (n=10) mice. Both genotypes started off at around 50% correct trials (black dotted line) in Session 1, which implies the mice had no bias for either image at the start of the task. There was no significant difference between genotypes in learning during the pairwise discrimination (two-way ANOVA, interaction GENOTYPE X SESSION F (1, 16) = 1.144, p=0.3007). **f** There was no significant difference in the number of days to reach criteria between genotypes (p=0.2093, t=1.162 df=16, two-tailed Student’s t-test). Values are plotted as mean ± SEM.

The memory impairments identified in *Kptn^−/−^* mice with the SOR and Barnes maze assays indicate a likely deficit in hippocampus-dependent memory retention. To test whether the *Kptn* KO mice also display impairments in tasks less reliant on hippocampal function, we used pairwise discrimination (PD), a visual operant conditioning task using a touchscreen platform^49–52^. During the PD task, both genotypes selected the correct image approximately 50% of the time in Session 1 (Fig. 2e, *Kptn^−/−^*, 45%; *Kptn^+/+^*, 53%), indicating that the mice had no innate bias for either of the images at the start of the task. The mean number of days to reach criteria (80% correct trials for two consecutive days^50^) was not significantly different between genotypes, and *Kptn^−/−^* mice performed no worse than *Kptn^+/+^*mice (*Kptn^−/−^*, 11.25 days; *Kptn^+/+^*, 14.5 days) (Fig. 2e). We performed a single reversal trial to confirm that the mice were capable of discriminating between images. As expected, the percentage of correct trials dropped significantly below 50% for both genotypes when the CS+ (rewarded) and CS- (non-rewarded) images were swapped (*Kptn^−/−^*, 24%; *Kptn^+/+^*, 19.2%), indicating a strong association with the original CS+ image (data not shown). These data indicate that *Kptn^−/−^* mice perform at least as well as wild-type animals in the PD task. The results of our three independent memory tests suggest that *Kptn* KO mice experience selective deficits, with at least some memory sparing.

### Cognitive sparing in patients with *KTPN*-related disorder

Our observation that certain cognitive traits were spared in our mouse model led us to investigate whether a similar outcome was detectable in patients with KRD. Six Amish individuals with KRD (aged 11-29, 3 female and 3 male individuals, Supplementary Fig. 2) and six age-matched Amish control individuals were psychometrically assessed for verbal comprehension, perceptual reasoning, processing speed, working memory, full scale IQ, list learning and narrative memory (see Methods). We detected significant impairment of cognitive function in all individuals with KRD, with all scores within the impaired range (z score < −2) compared to matched controls, except the relative sparing of narrative memory function (Fig. 3). As narrative memory can be considered to have less of a hippocampal contribution^53^, these psychometric test findings are consistent with the results found in our mouse model. This cross-species comparison highlights the concordance of our mouse model with the human disorder at a cognitive level.

**FIGURE 3.**
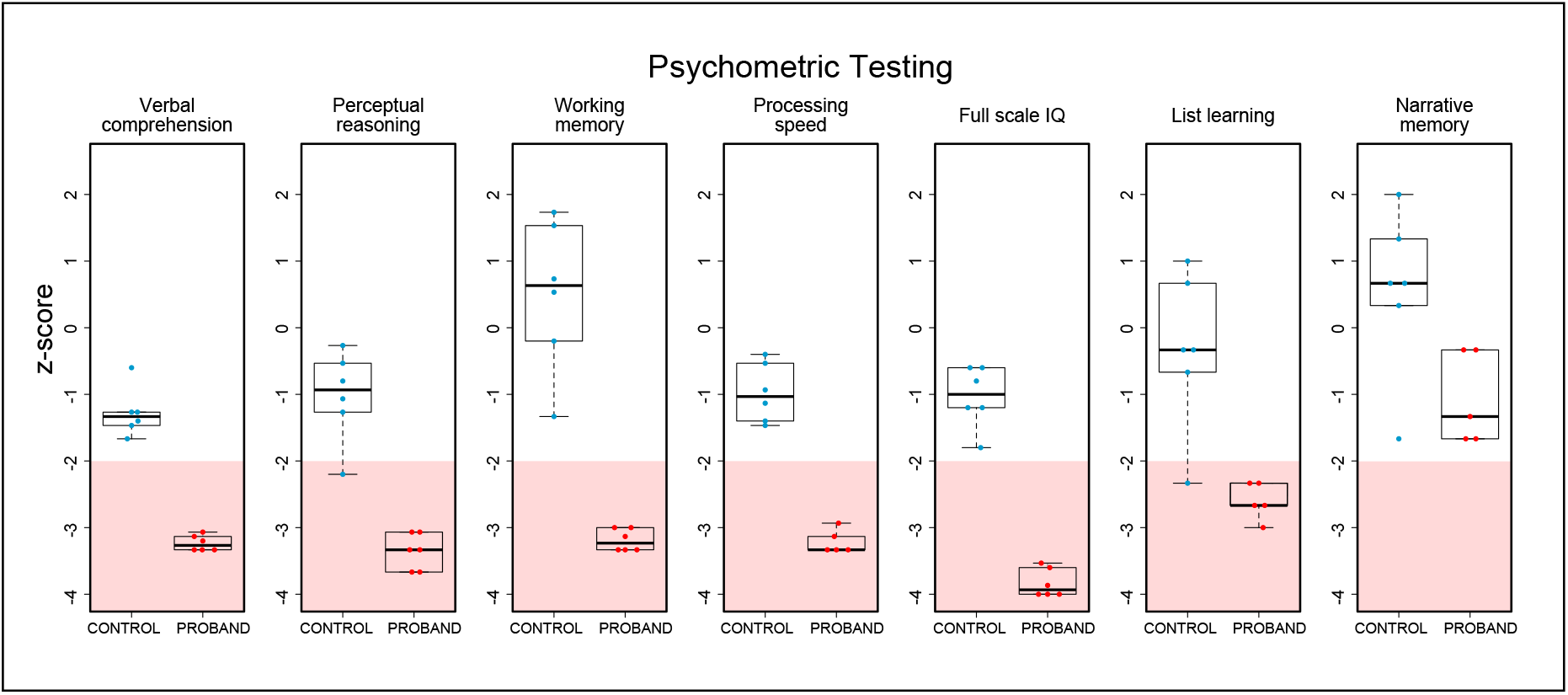
Psychometric testing of Amish individuals with KRD reveals cognitive sparing. Six Amish individuals between the ages of 11 and 29 with KRD (3 female, 3 male) and age-matched population controls were psychometrically assessed (see methods) to measure cognitive performance in four domains including verbal comprehension, perceptual reasoning, processing speed and working memory indices, that can be combined to generate a full-scale intelligence quotient score. Auditory-verbal recall performance was also assessed by list learning and story memory tests. KRD patients were consistently in the impaired range (z-score < 2, pink shading) except in the narrative memory assessment, probands all fall outside the impaired range.

### *Kptn* deficiency is associated with severe and progressive macrocephaly in mice

Macrocephaly and frontal bossing are frequently reported features of KRD in humans. We therefore investigated whether the *Kptn^−/−^* mice also replicate this phenotype. Microcomputed tomography (μCT), magnetic resonance imaging (MRI) and histological morphometrics were used to quantify changes in skull and brain volume, shape, and cellular distribution^54^.

Applying μCT-based cephalometric analysis to examine skull size, we detected an increase in the height, length, and width of the brain cavity of the skulls of adult *Kptn^−/−^* mice (Fig. 4a-d, Supplementary Table 1, Supplementary Fig. 3). The height of the skull at the midline is consistently elevated throughout the rostro-caudal extent of the brain cavity (5.2% at Bregma/L1-L7, 4.7% at Lambda/L2-L5, p<0.01) as seen in a sagittal resection of representative 3D volumes (Fig. 4b). In coronal resections, the height changes are due primarily to an increase in the dorsal curvature of the frontal and parietal bones (Fig. 4c,d), rather than an increase in the dorsoventral length of the lateral skull wall. This dorsal bulging is consistent with the changes observed in frontal bossing in humans, where the frontal bone may expand to accommodate pathological brain overgrowth^55,56^. Applying MRI tensor-based morphometry followed by voxel-based quantification of brain volume^57^, we detected a 7.7%/9.2% increase (female/male, p<0.001) in mean intracranial volume in *Kptn^−/−^*mice at 16 weeks when compared to wild-type controls (Fig. 4e). Together, these bone and soft-tissue findings indicate that *Kptn^−/−^* mice exhibit both skull shape abnormalities, including overgrowth and deformation, and megalencephaly. These are consistent with the macrocephaly and frontal bossing seen in KRD.

**FIGURE 4.**
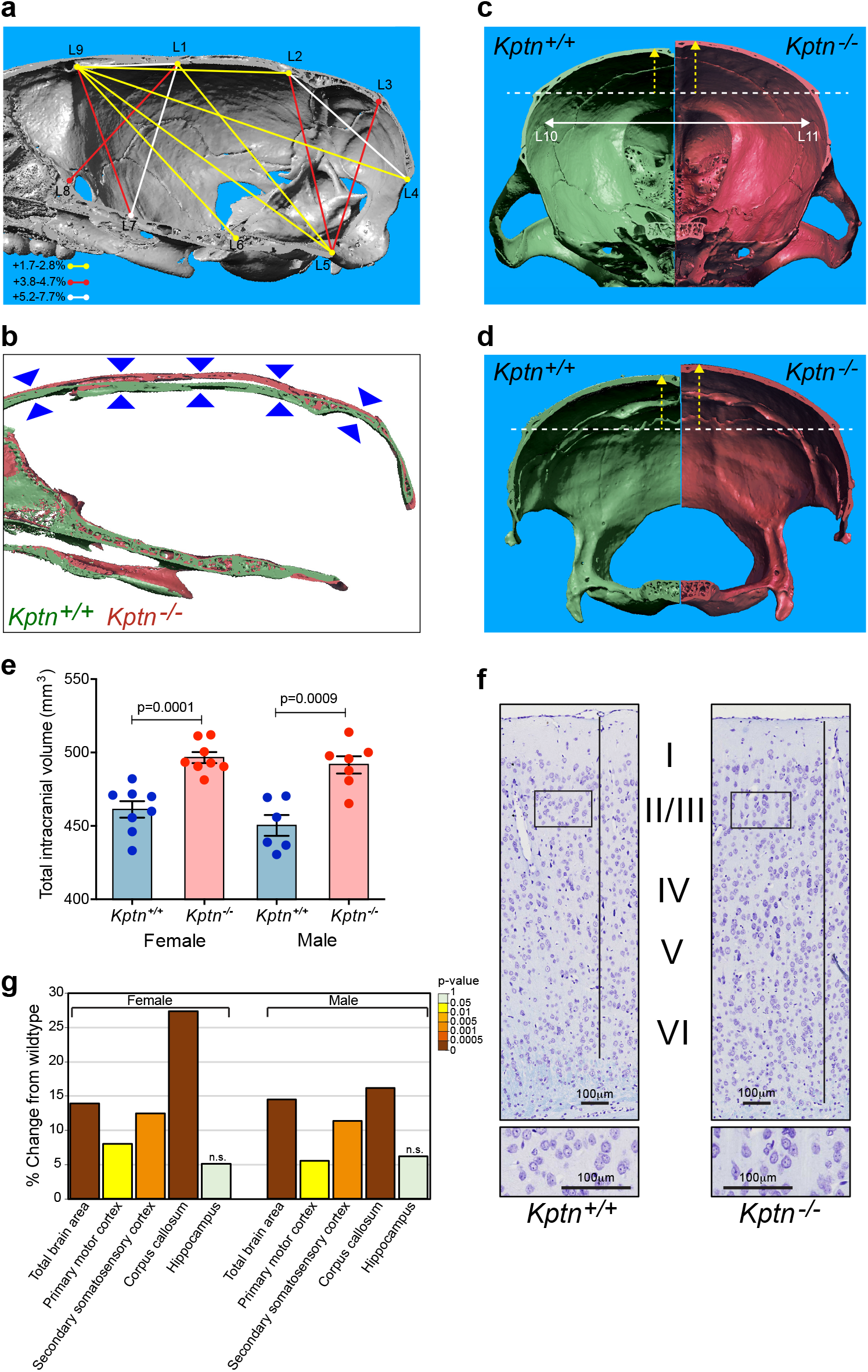
*Kptn*^−/−^ mouse model exhibits skull and brain overgrowth. Micro-computed X-ray tomography reconstructions were collected and analysed on male *Kptn*^−/−^ mice and *Kptn*^+/+^ controls (n=5 each). **a** Significant increases in inter-landmark distances were observed in height, length, and width of the brain cavity of *Kptn*^−/−^ animals (1.7-7.7% increase in mean linear distances, p<0.05, two-tailed Student’s t-test). **b** Sagittal sections of 3D reconstructions illustrate the increased skull height along the full rostro-caudal extent of the brain cavity (arrowheads). **c,d** Rostral and caudal-facing reconstructions of coronal sections from representative individual 3D reconstructions highlight changes in the dorsal curvature (yellow arrows) of frontal and parietal bones in *Kptn*^−/−^ mice (left, green = *Kptn*^+/+^, right, red = *Kptn*^−/−^), and the width of the brain cavity (white arrow, +2.56%, p= 0.0356). **e.** Volumetric measurements from MRI of female and male *Kptn*^−/−^ mutants and *Kptn*^+/+^ controls detect 7.7%-9.2% (F/M) increases in total intracranial volume at 16 weeks of age (p<0.001, two-tailed Student’s t-tests). Values are plotted as mean ± SEM. **f** Representative sections from *Kptn*^+/+^ and *Kptn*^−/−^ cortices stained with cresyl violet. **g** Morphometric analyses of histological sections (as in **f**) of *Kptn*^−/−^ mutant and *Kptn*^+/+^ controls plotted as percentage difference of *Kptn*^−/−^ from wildtype mean, indicating significant increases in total brain area (shown for section 1), cortical (shown for section 1) and corpus callosum thicknesses (section 2) (p<0.002 in most measurements as indicated, and Supplementary Table 2, two-tailed Student’s t-tests). The hippocampus was not significantly (n.s.) changed in size (p>0.2).

We next characterised the megalencephaly using previously published histomorphometric methods^58,59^, quantifying neuroanatomical features across the same three coronal brain regions in all subjects. Substantial brain size anomalies were identified in both adult female and male *Kptn^−/−^* mice (16 weeks) when compared to matched wildtypes, with 34 out of 78 parameters affected (Fig. 4f, Supplementary Fig. 4a-c, Supplementary Table 2a-c). The total brain area in both sexes was significantly increased across the three coronal sections (+7.9-14.4%, p≤0.0007), concomitant with enlarged cortices in all areas measured, and enlarged corpus callosum structures (Fig. 4f,g). The lateral ventricles were the only brain regions exhibiting a decreased size (section 1: −36%, p=0.05; section 2: −54%, p=0.02), likely indicating a compensatory reaction to increased brain volume constrained within the skull. Interestingly, we detected no significant difference in hippocampus-related size parameters (Fig. 4g, Supplementary Fig. 4a-c, Supplementary Table 2a-c). The histological findings are consistent between sexes (Supplementary Fig. 4a-c, Supplementary Table 2a-c). Of note, we find that the brain enlargement is accompanied by increased cell number rather than a change in cell body size (Supplementary Table 2d), as has been observed in some models of mTOR related macrocephaly^60^.

To better understand the developmental trajectory of brain overgrowth in our mouse model, we compared histomorphology at birth (Postnatal day P0), P20, and 16 weeks of age. P0 allows us to assess whether *Kptn* mutant animals are born with enlarged brains, or conversely overgrow postnatally. P20 represents the stage at which wild-type mice reach 90-95% of adult brain weight^61,62^ and thus allows us to distinguish between an increased growth rate and an extended brain growth period. In the former, we would anticipate a larger brain by the end of normal brain growth (P20), while the latter would lead to later appearance of overgrowth (16 weeks). Using histological methods described above, we detected no phenotypic differences at P0 in the vast majority of parameters (51 out of 53) including the total brain area (Supplementary Fig. 4e, Supplementary Table 2e,f), indicating that the megalencephaly phenotype is associated with postnatal developmental processes. Interestingly, two parameters were altered at P0, the total area of the hippocampus is reduced by 11% in the *Kptn^−/−^* mice (*p*=0.04), and the internal capsule increased by 9% (*p*=0.03), suggesting that some morphological anomalies originate from prenatal stages but these are highly tissue restricted (Supplementary Fig. 4e, Supplementary Table 2e,f).

A similar analysis on P20 mice again identifies very few statistically significant changes in regional brain size (Supplementary Fig. 4d, Supplementary Table 2g), despite a broad upward trend. Consistent with all stages examined, the ventricles of P20 *Kptn^−/−^* mice are reduced in size, supporting the idea that there is ongoing, but mild overgrowth at early stages that continues into adulthood, eventually resulting in severe megalencephaly. It is striking that *Kptn^−/−^* mice lack significant overgrowth at P20, when wild-type mice will have reached the end of the normal brain growth. The progressive nature of the disorder is well-illustrated when we plot measurements of total brain area, motor cortex, and hippocampus with time (Fig. 5a). We note that the underdevelopment of the hippocampus at P0 recovers with age, but never reaches statistically significant overgrowth during the life of the mouse. Taken together with the significant brain size changes in adult *Kptn^−/−^* mice, these data indicate that the megalencephaly seen in the *Kptn* model is postnatal, progressive, and results from an extended period of brain growth.

**FIGURE 5.**
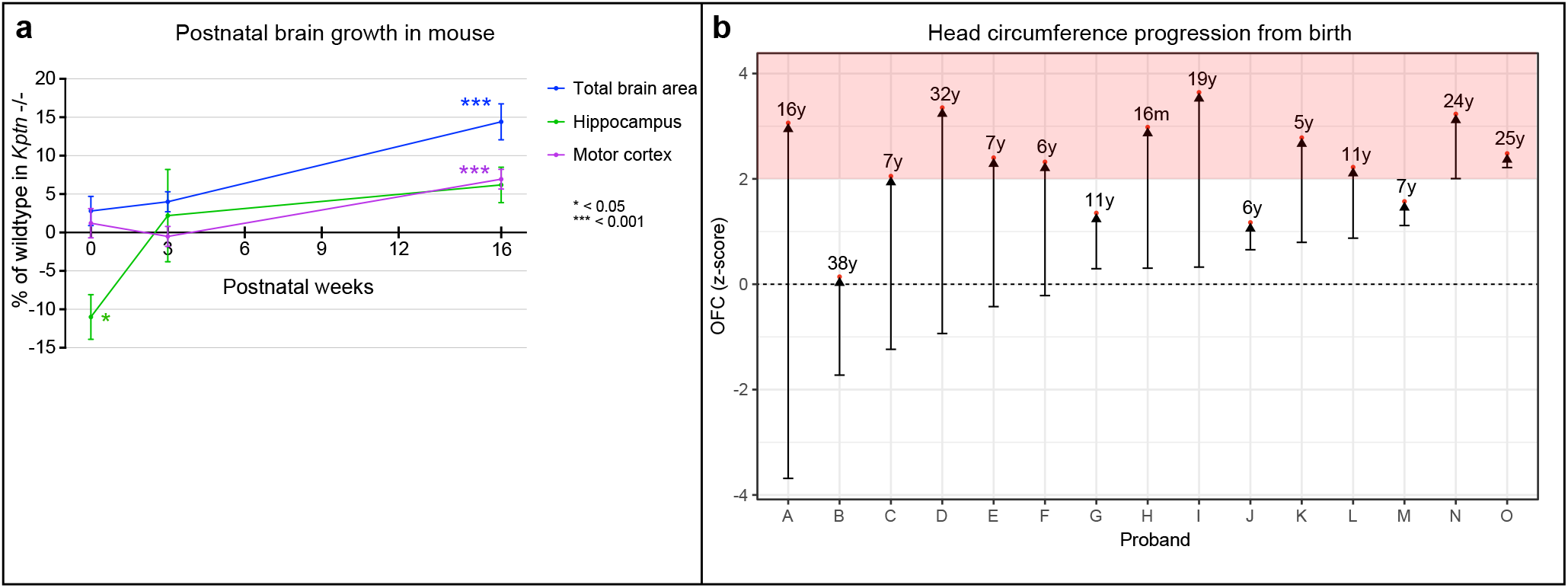
Mouse KRD model and human KRD probands display postnatal brain overgrowth. **a** Percentage mean change of total brain area, motor cortex and hippocampus in *Kptn*^−/−^ mutant mice at birth, 3 weeks, and 16 weeks postnatally. Significant overgrowth is detected only in 16 week animals (p<0.001, two-tailed Student’s t-test). **b** Occipital Frontal Circumference (OFC) measurements from fifteen individuals with KRD from birth (lower bars) to age at last assessment (red dots) indicate that overgrowth occurs primarily postnatally. Pink shading in **b** indicates the macrocephalic range, with SD > 2.

### Macrocephaly in *KPTN*-related disorder is postnatal and progressive

We set out to clarify whether the macrocephaly seen in individuals with KRD is also postnatal and progressive. This could provide opportunities to intervene in the disorder progression after birth, thus increasing the feasibility of any proposed treatment. We identified a total of 35 patients with a confirmed molecular diagnosis of KRD (Supplementary Table 3), including 20 new Amish and non-Amish individuals through our international collaborators, revisited the 9 affected individuals originally described from the Amish community, and reviewed OFC data for 6 from previously published studies^28–30,32^. Fig. 5b shows the OFC trajectories of 15 KRD patients from birth to their most recent available assessment. The majority of KRD patients are born with OFCs within the normal range, and progress with age to become macrocephalic (>2 SD, pink zone in Fig. 5b), with the most rapid increase occurring within the first 2 years of life (representative OFC graph in Supplementary Fig 5). These data demonstrate further consistency between the human disorder and our mouse model, which both experience progressive postnatal overgrowth.

### Kptn regulates mTOR signalling *in vivo*

KPTN functions in a protein complex (KICSTOR) negatively regulating mTOR activity in response to nutrient availability^19,33^. Given the known contribution of mTOR pathway members in numerous brain overgrowth disorders in humans^9,63–65^, this provides a plausible mechanism for KRD overgrowth. To examine whether KPTN modulates mTORC1 signalling in our *in vivo* model, we performed Western blot assays on brain tissue from *Kptn^−/−^* and *Kptn^+/+^* mice assessing ribosomal protein S6 (RPS6), a known downstream target of the mTORC1 pathway whose phosphorylation state is strongly linked to pathway activation^66,67^. Our results in both whole brain and hippocampus of adult animals reveal a significant increase in mTOR pathway activity in *Kptn*^−/−^ mice, as indicated by increased phosphorylation of RPS6 (p < 0.05; Fig. 6a). Importantly, we find mTOR activation is present at P21 (p=0.0308, Fig. 6a), the stage at which the *Kptn*^−/−^ model is experiencing brain overgrowth. Immunostaining for phosphorylated RPS6 (p-RPS6) in histological sections of brain tissue highlights the widespread cortical and hippocampal activation of mTOR activity in *Kptn*^−/−^ mice (Fig. 6b). To confirm that the increase in p-RPS6 caused by loss of *Kptn* is mTOR dependent, we treated a cohort of mice with rapamycin, an mTOR pathway inhibitor^68–70^. We find that the excess p-RPS6 signal seen in *Kptn*^−/−^ animals is significantly reduced after 3 days of treatment, confirming the presence of hyperactive mTOR signalling in our model (p=0.00188, Fig. 6c). These data are consistent with the proposed role of Kptn as a negative regulator of mTORC1 activity and provides *in vivo* evidence of a role for mTOR signalling in KRD.

**FIGURE 6.**
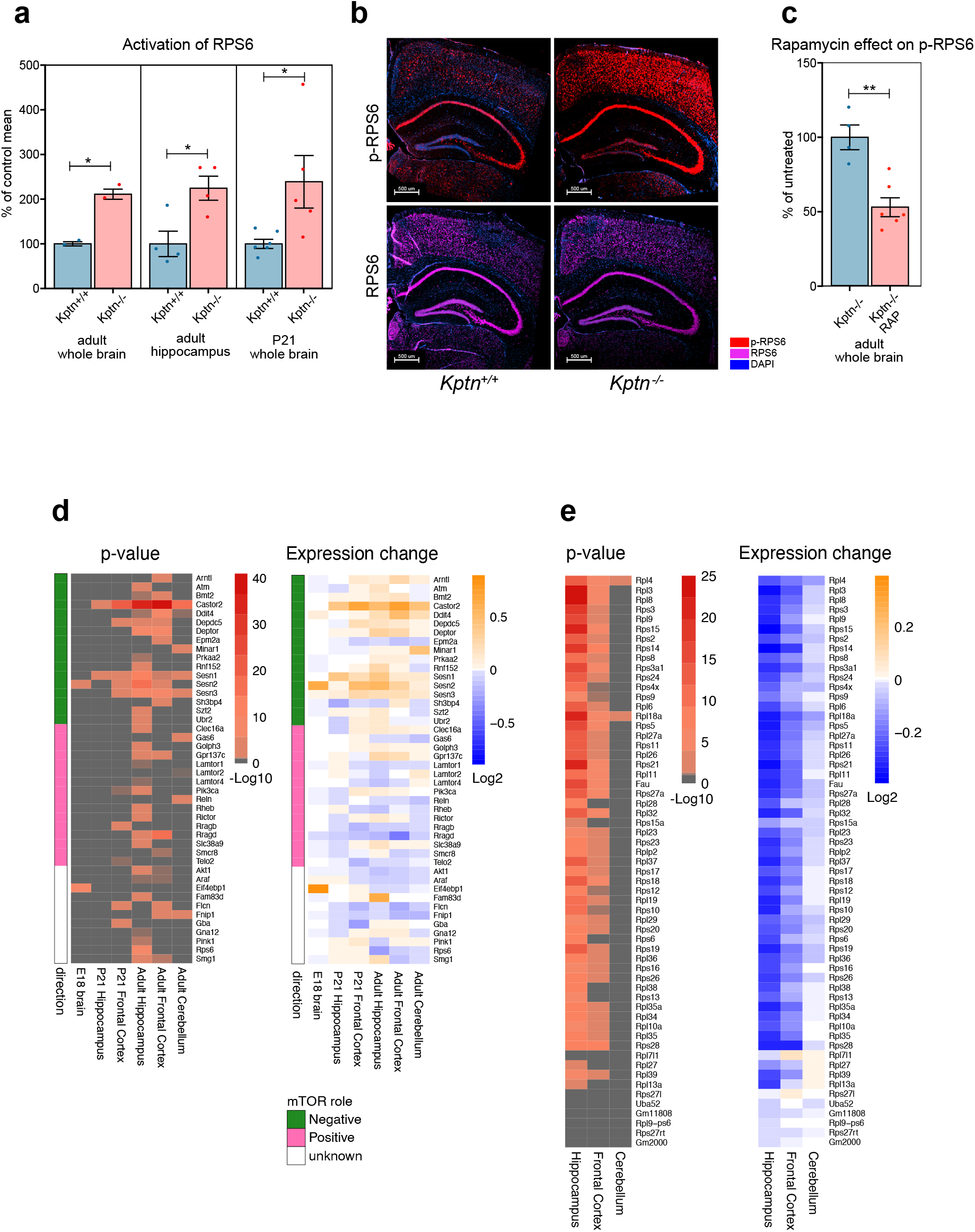
Kptn loss-of-function induces changes in mTOR signaling corrected by rapamycin. **a** Quantification of the phosphorylation of Ribosomal Protein S6 (RPS6) in wild type controls (*Kptn^+/+^*) and mutant animals (*Kptn*^−/−^) indicate significant increases in mTOR signaling in adult whole brain (p=0.0119, n=2 per genotype), hippocampus (p=0.0189, n=4 per genotype), and in juvenile mice at P21 (p=0.0308, *Kptn*^+/+^, n=6; *Kptn*^−/−^, n=5), measured as the ratio of phosphorylated RPS6 (p-RPS6) to total RPS6 signal (units as % of control mean). All p-values are from two-tailed Student’s t-tests). **b** Immunostaining for RPS6 (purple) and its phosphorylated form (p-RPS6, red) reveal widespread increases in cortical and hippocampal activation of the mTOR pathway in *Kptn*^−/−^ mice (nuclei stained with DAPI, blue). **c** Quantification of the phosphorylation of RPS6 in *Kptn*^−/−^ animals after 3 days of treatment with vehicle (*Kptn*^−/−^) or rapamycin (*Kptn*^−/−^ RAP) indicates a significant reduction in p-RPS6 levels upon treatment (p=0.00188, *Kptn*^+/+^ n=4; *Kptn*^−/−^ n=6, two-tailed Student’s t-test). Values are plotted as mean ± SEM. **d,e** transcriptional changes to mTOR pathway components (**d**) and downstream ribosomal gene network (**e**) in *Kptn*^−/−^ mice at embryonic and postnatal stages as indicated. Statistically significant (α<0.05) Log_2_-fold changes in expression are indicated by non-grey adjusted p-value heatmap cells. Role in mTOR pathway is indicated as green/pink/white colour scheme in **d**.

### Transcriptional changes in *Kptn* animals and human neural cells highlight mTOR pathway dysregulation

While few transcriptional readouts have been described for mTOR activity, the tight regulation of this critical metabolic pathway involves extensive use of feedback loops to maintain homeostasis^71^. We hypothesized that the loss of negative mTOR regulation in our *Kptn*^−/−^ model would likely result in detectable changes to the expression of these mTOR pathway members, and genes involved in the downstream cellular processes. We performed RNA-Seq on dissected P21 and adult brain tissues (prefrontal cortex, hippocampus, and cerebellum), comparing gene expression between *Kptn*^−/−^ and *Kptn^+/+^* animals. Extensive dysregulation of both positive and negative regulators of the mTOR pathway were observed (Fig. 6d). Interestingly, adult hippocampus and cortex show strong dysregulation, concordant with the immunostaining results in adult tissue. A notable observation is the enrichment in *upregulation* of negative mTOR regulators (Fig. 6d), likely reflecting a strong negative feedback response in an attempt to attenuate the hyperactivated mTOR signalling resulting from loss of Kptn protein. Numerous genes involved in protein synthesis, including most ribosomal protein genes, are dysregulated in the *Kptn*^−/−^ mice (Fig. 6e, Supplementary Table 4a). These two observations are supported by the statistically significant enrichment of differentially expressed genes related to these processes, when assessed by unbiased Gene Ontology enrichment^72^ (Supplementary File 1a). When we test specifically for dysregulation of mTOR pathway genes, 5 out of 6 tissues examined show a significant enrichment (hypergeometric test p<0.05, Supplementary File 1b). These data are concordant with the known regulation of ribosomal protein genes by mTOR and its critical role in controlling protein synthesis^73–75^.

To confirm the relevance of our mouse findings to human biology, we generated human induced pluripotent stem cell (iPSC) models of KRD. We selected a guide RNA (gRNA) targeting exon 8 of the human *KPTN* gene, directly between two previously identified causal variants, aiming to largely recapitulate the known nonsense variant (p.Ser259*)^28^. Using CAS9 riboprotein electroporation in the KOLF2C1 cell line^76^, we isolated clones carrying frameshifting indels in exon 8 of *KPTN* (Supplementary Fig. 1a). In order to examine the effect of *KPTN* LoF in a biologically relevant cell type, we differentiated our iPSC models into PAX6-positive cortical neural precursor cells (NPCs) using established methods^77^. The functional consequence of the frameshifting variants was characterised by RNA-Seq, which reveals a dose-dependent reduction in detectable *KPTN* transcript reads in heterozygous (41% reduction) and homozygous (89% reduction) mutant NPC lines (Supplementary Fig. 1c), likely reflecting robust nonsense mediated decay of the mutant transcripts.

Transcriptomic analysis of these human brain stem cell models of KRD and isogenic wildtype controls (Supplementary Table 4b) reveals significant differential expression of numerous genes involved in mTOR signalling, and nearly complete dysregulation of the ribosomal protein gene network (Fig. 7a,b). The TSC proteins, important inhibitors of the mTOR pathway, are both downregulated in *KPTN*^−/−^ NPCs. LoF variants in TSCs are associated with tuberous sclerosis complex, a multi-system tissue overgrowth syndrome affecting brain, skin, kidney, heart, and lung^78^. We find that 36% of genes dysregulated in TSC2^−/−^ neurons^79^ are also differentially expressed in our *KPTN*^−/−^ NPCs (301 out of 834 expressed in NPCs, 1.35 fold over-enriched, hypergeometric test p=4.3×10^−10^), thus identifying a common signal in neural cell types driven by convergent disease mechanisms affecting the mTOR pathway.

**FIGURE 7.**
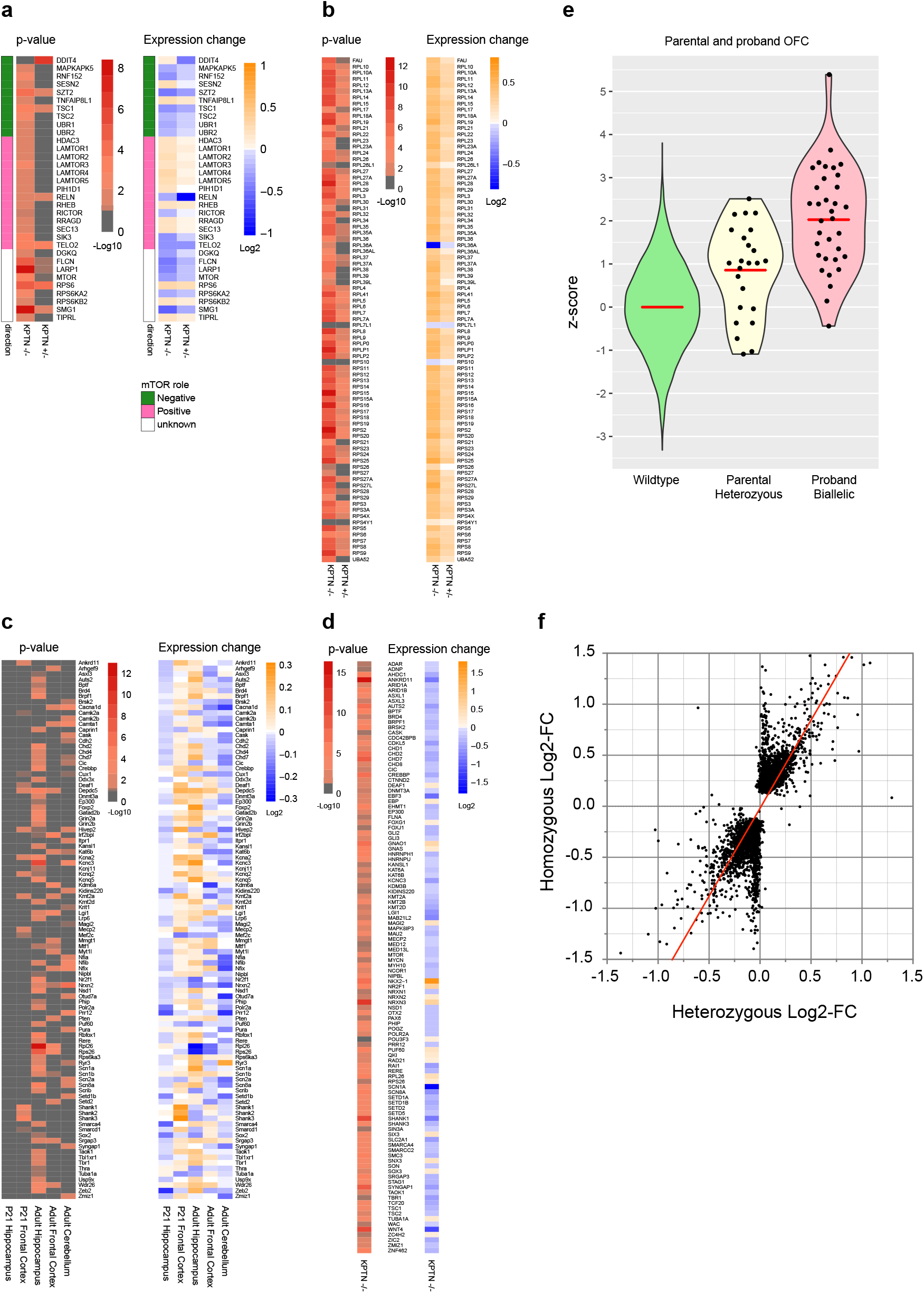
Human neural stem cell model of KRD has mTOR pathway and developmental disorder gene dysregulation. **a, b** KPTN LoF NPC models display changes in gene expression of mTOR pathway components (**a**) and downstream ribosomal gene network (**b**). Heterozygous *KPTN* LoF cells (*KPTN^+/−^*) have an intermediate mTOR pathway phenotype consistent with dosage sensitivity to loss of *KPTN* expression. **c** Mouse brain and **d** human neural cell models of KRD show significant dysregulation of dominant haploinsufficient developmental disorder associated genes (statistically significant Log_2_-fold changes in expression are indicated by non-grey adjusted p-value heatmap cells). **e** OFC measurements in heterozygous parents (Parental) of KRD probands reveal a significant increase in mean head circumference (mean z-score=0.856, p=0.000028, one sample, 2-sided z-test against wildtype distribution, n=24) compared to wildtype OFC distribution, and an intermediate effect relative to that observed in affected individuals with biallelic *KPTN* variants (Proband, mean z-score=2.027, p<0.00001, one sample, 2-sided z-test against wild type distribution, n=35). **f** RNA-Seq on *KPTN* LoF NPC models indicates a strong correlation between the gene dysregulation in homozygous and heterozygous KPTN cell lines (Pearson r=0.759, p<0.0001), consistent with a dose-dependent effect on pathway activity.

The Connectivity Map^80^ (CMAP), a large-scale collection of transcriptional response profiles after compound exposure (20,000 different compounds and drugs) or gene knockdown across numerous cell lines, uses negative correlation between drug-disease profile pairs to identify potential therapeutics to reverse disease states. We queried the CMAP database with the transcriptional response signature of our *KPTN*^−/−^ NPCs. We find that torin-2^81^, a known mTOR inhibitor, scores as the most strongly negatively correlated profile (global correlation score of −96.28 across all cell lines, Supplementary Fig. 6a), and is therefore considered the most promising treatment for the queried disease-profile. Interestingly, in HCC515 cells five of the top 32 anti-correlated signatures are mTOR inhibitors (torin-2, QL-X-138, torin-1, sirolimus, and WYE-354) with scores between −99.56 and −93.23 (Supplementary Fig. 6b). This strong anticorrelation with mTOR inhibitor responses, detected in non-neural cancer lines, suggests that the effect of loss of KPTN protein on the mTOR pathway is robust and specific.

### Increases in radial glial and intermediate precursor marker expression

It has been shown that the mTOR pathway is both active in and critical for the development and differentiation of neural stem cells in both mouse and humans^22,23,82^. Pathological increases in mTOR signalling, seen in tuberous sclerosis complex and somatic *MTOR* variants linked to hemimegancephaly^25,26^, point to a significant input of the mTOR pathway in the control of neural stem cell expansion and differentiation. To understand how the loss of KPTN protein is leading to hypercellularity in KRD, we examined the expression of markers of neural stem cells and radial glia, and those of proliferating neuronal progenitors in our mouse model. We find significant increases in levels of radial glial markers (e.g. *Nestin, Vimentin, GLAST/Slc3a1, and Thrsp*) and intermediate progenitor markers (e.g. *Tbr1, Eomes/Tbr2, Neurod2, Btg2*)^83–87^ in the mouse brain at both P21 and adult stages (Supplementary Table 5). These striking increases in gene expression support a model in which the persistence, beyond P21, of excessive neural stem cells and cycling neural progenitors contribute supernumerary neuronal cells, resulting in significant pathological brain overgrowth. These data provide an important starting point for further work on the cellular basis of megalencephaly in KRD.

### Epilepsy gene networks in *KPTN*-related disorder

Around half (47%) of KRD patients experience seizures^28–32^. Although we did not observe any clinical seizures during the maintenance and testing of *Kptn*^−/−^ mice, the absence of detectable spontaneous seizures in our model is not totally unexpected. Other mouse models of human mTOR-associated epilepsies such as *SZT2*^33^ (a KPTN binding partner and mTOR regulator) show a reduced threshold for seizures after pentylenetetrazole and electroconvulsive treatment, while lacking the spontaneous seizures observed in the corresponding human disorder^88,89^. Transcriptomic analysis of our KRD models, however, reveals the dysregulation of numerous known epilepsy associated genes^90,91^ (syndromic and non-syndromic) in both mouse brain (290 out of 844 expressed, Supplementary Fig. 7a) and cortical NPCs (281 out of 807 expressed, Supplementary Fig. 7b). Among the differentially expressed seizure genes, around half of the mouse genes (n= 122) and a third of the human genes (n= 97) are concordant between direction of change and known mechanism of disease (e.g. downregulated expression of a LoF mechanism disease-associated gene). There is a significant overlap in differentially expressed seizure genes between mouse and human KRD models (107 genes in common, 1.5 fold over-enriched, hypergeometric test p=3.6×10^−7^, Supplementary Fig. 7c). Of particular interest are genes known to be dosage sensitive and are therefore most likely to contribute to seizure predisposition. For example, *SCN1A, SCN8A, SLC2A1, SYNGAP1, SHANK1*, and *LGI1* all show more than 30% reduction in expression in *KPTN*^−/−^ NPCs. *KCNQ3*, a potassium channel protein found mutated in neonatal seizures, shows a 25% increase in expression in these cells, consistent with the suspected altered-function mechanism in patients^92,93^. Of others known to operate by an altered or gain-of-function mechanism, *Sik1* is of note, with a 90% increase in expression in the prefrontal cortex of *Kptn*^−/−^ mice (p-adj = 0.0012). Patients carrying *SIK1* missense mutations experience severe developmental epilepsy, with mutations altering the protein’s ability to correctly regulate MEF2C activity in neurons^94,95^. Taken together, these data suggest that KPTN LoF is resulting in a dysregulation of numerous genes that control electrophysiological homeostasis in brain tissue, leading to a susceptibility to seizures in affected individuals.

### Dosage sensitive gene networks are disrupted in KRD models

The regulation of mTOR signalling is critical to properly regulated growth and differentiation during development^11,96^. The loss of an important negative regulator of this pathway during development would therefore be expected to disrupt important gene networks that regulate these processes. Indeed, we observed that numerous genes causally linked to other neurodevelopmental disorders are differentially expressed in our KRD models. Haploinsufficient genes (dosage sensitive disease-associated genes, Fig. 7c,d), whose phenotype appears in heterozygous loss-of-function, are key regulators of brain development and normal brain function^97^. We find a surprisingly consistent downregulation of these (82% of 111 genes, Fig. 7d) in *KPTN*^−/−^ human cortical neural precursors (p<0.00001, Fisher’s exact test). This includes a more than 25% reduction in expression of important chromatin modifying proteins and transcriptional regulators *AUTS2, AHDC1, ANKRD11, CHD7, CREBBP, CHD2, EBF3, KMT2D, TBR1, TCF20, SETD1B* and *ZIC2*. Individual disruptive variants in each of these genes results in severe neurodevelopmental deficits in KRD-relevant phenotypic areas (Supplementary Table 6). Substantial changes in the expression of these proteins indicates a significant impact of *KPTN* gene function on a broad network of developmentally important genes, whose reduced expression has a known impact on cognitive function.

### Human brain growth is sensitive to KPTN dosage

During clinical data collection from Amish individuals affected by KRD, it was observed that several of their parents presented with larger than average head size. As these parents are heterozygous carriers of pathogenic *KPTN* gene variants, we carried out a systematic analysis of OFC in 24 parents of identified KRD cases. We observed a significant increase in head size (mean z-score =0.856, p=0.000028) in carriers compared to the normal population distribution, though infrequently in the macrocephalic range (4 out of 24 had an OFC >2 SD, Fig. 7e). This surprising observation suggests that while KRD is a recessive condition, *KPTN* alleles have an additive effect on head size, with reduced KPTN dosage in heterozygous carriers leading to head overgrowth, possibly due to the same mechanisms of mTOR hyperactivation. To test this directly, we differentiated heterozygous *KPTN* LoF iPSCs into cortical neural precursors. RNA-Seq on these cells reveals a strong correlation between the gene dysregulation in homozygous and heterozygous *KPTN* lines (Pearson r=0.759, p<0.0001, Fig. 7e). We find an intermediate mTOR pathway dysregulation in the heterozygous cells, and widespread disruption to the ribosomal protein gene network, consistent with that observed in the homozygous *KPTN* lines (Fig. 7a,b, Supplementary Table 4b). These data strongly suggest that human head size is sensitive to *KPTN* gene dosage, likely through its action within the mTOR pathway. Although none of the heterozygous carrier parents had any reported cognitive impairment or history of seizures, further studies will be required to understand whether the growth effect is accompanied by any changes in neurological function.

### Conclusions

We have generated and characterised two informative models of *KPTN*-related disorder, a recent addition to the growing family of mTORopathies. Through behavioural, cognitive, morphological, and molecular investigations we have demonstrated robust concordance between the *Kptn^−/−^* mouse and numerous aspects of human KRD. Our detailed characterisation of the progressive nature of brain overgrowth in KRD provides critical insight into disease progression. Through animal and cellular modelling, we have strengthened the link between KPTN and mTOR signalling, and by inference, the role of aberrant mTOR signalling in KRD.

Our pre-clinical therapeutic investigation indicates that mTOR dysregulation in KRD is rapamycin sensitive, providing a pathway to possible clinical intervention. The relationship between megalencephaly and cognitive dysfunction in KRD is unclear. The brain overgrowth phenotype could be the primary cause of the cognitive deficits observed in KRD. Alternatively, the neurological phenotypes could be caused by the persistent deregulation of mTOR signalling we observe in adult *Kptn^−/−^*mice, given the known role of mTOR in cognition and memory formation^14^. If so, the morphological changes in KRD could simply reflect the effects of *KPTN* variants on brain growth via overactive mTOR signalling. Indeed, the neurological features of KRD are strikingly similar to Smith-Kingsmore syndrome caused by germline gain-of-function variants in *MTOR*^27^. As mTOR inhibitors are generally well tolerated^98,99^, the path to clinical intervention in KRD may be tantalisingly short, and follow-up experiments using our mouse and cellular models should provide critical information to shape clinical trials.

Our study highlights the strength of combining clinical investigation with the ongoing characterisation of disease models, allowing a more thorough description of both human and model disease pathophysiology. Using our mouse findings in cognitive function, we were able to subsequently identify areas of cognitive sparing in affected individuals. Our analyses of mouse model head morphology revealed a concordant widespread overgrowth in both brain tissue and skull bones. Through a careful examination of the developmental trajectory of brain overgrowth in our mouse model, we were able to confirm the progressive nature of KRD macrocephaly. Conversely, we were able to take clinical observations from apparently unaffected parents and confirm in cellular models that the heterozygous carriers of pathogenic *KPTN* LoF variants likely experience mTOR dysregulation, contributing to a significant increase in head circumference. While the role of somatic and germline mutations affecting mTOR signalling in pathological brain overgrowth is well established^25,100,101^, we were surprised to find such a strong, yet previously unrecognised phenotype effect of brain overgrowth in heterozygotes. There is increasing evidence of a role for recessive disease alleles in causing incompletely penetrant phenotypes in the general population, as the rapidly growing sequencing efforts provide the necessary power to detect these effects^35–38^. While *KPTN* heterozygotes are not known to experience any neurological consequences, constraint metrics such as LOEUF^102^ indicate that *KPTN* LoF alleles may be experiencing purifying selection beyond what would be expected from a purely autosomal recessive disorder gene (observed/expected LoF=0.5, LOEUF=0.81). Further investigation into the phenotypic consequences of carrier state will be needed to clarify this apparently benign effect on head size, and indeed whether natural variation in other mTOR pathway genes is detectably contributing to differences in brain growth (as suggested in Reijnders *et al.*, 2017).

By exploring the extent and character of transcriptional dysregulation in our KRD models, we identified perturbations to numerous mTOR pathway members, seizure-related genes, and transcriptional regulators essential to normal brain development and function. The persistence of progenitor marker genes (for both radial glia and intermediate neuronal precursors) beyond their normal period of expression supports a model in which chronic hyperactivation of the mTOR pathway during postnatal brain development may result in the pathological persistence of proliferating progenitor cells, and the supernumerary differentiation of neural cell types beyond P21. These dysregulated postnatal developmental processes likely result in significant progressive brain overgrowth. The structural changes are likely compounded by chronic mTOR hyperactivation in the adult brain, leading to significant neurocognitive phenotypes in our models and in KRD in humans.

Our multimodal analysis of mouse and cellular models of KRD provides a deeper insight into the underlying mechanisms of disease, and places KRD in the context of a family of disorders with a common underlying mechanism of mTOR dysregulation. Further work to refine our understanding of the timing and affected cell types during postnatal development will contribute to a more informed therapeutic strategy in treating this and other mTOR-related disorders. The promising treatment outcomes of related mTORopathy mouse models^98,104–106^ provide encouraging evidence supporting the aim of effective therapeutic intervention in KRD.

## Acknowledgements

First and foremost, we are grateful to the patients and their families for participating in this study. In particular we would like to thank the KPTN alliance and the Amish community for their support of the Windows of Hope project. This work was supported by core Wellcome funding to the Wellcome Sanger Institute (Grant reference numbers: 206194 and 108413/A/15/D), and Open Targets project OTAR2053. B.Y. is supported by funding from the Jerôme Lejeune Foundation, the French National Research Agency (ANR) and the National Institute of Health and Medical Research (INSERM). PBC, ELB and AHC are supported by a Javits Award (NINDS R37-NS125632). LER, ELB and AHC were also supported by the Newlife Foundation (16-17/12). We would like to thank all the staff of the Wellcome Sanger Research Support Facility for mouse husbandry and their contributions to rapamycin injection sessions, Hannah Wardle-Jones for expert colony management, and members of the Wellcome Sanger Mouse Genetics Programme for the initial mouse model observations. This study makes use of data generated by the DECIPHER community. A full list of centres who contributed to the generation of the data is available from https://deciphergenomics.org/about/stats and via email from. Funding for the DECIPHER project was provided by Wellcome. For the purpose of open access, the author has applied a CC BY public copyright licence to any Author Accepted Manuscript version arising from this submission.

## Data availability

The European Nucleotide Archive accession numbers for the RNA-seq sequences reported in this paper are as follows: Mouse-derived brain RNA-Seq samples: ERA941410-ERA941421, ERA941432-ERA941465, ERA2115013-ERA2115023, ERA2115035-ERA2115045 iPSC-derived NPC samples: ERS8443331, ERS8443330, ERS8443332, ERS8443333, ERS8443322, ERS8443323, ERS8443326, ERS8443327, ERS8443328, ERS8443329

## METHODS

### Mouse production

The mouse model was generated at the Wellcome Sanger Institute. The Kptn*^tm1a(EUCOMM)Wtsi^* mice were kept on a C57BL/6NTac background (Taconic Biosciences). The tm1a ‘knockout-first’ allele was generated by the insertion of an IRES:lacZ trapping cassette and a LoxP flanked promoter-driven neo cassette into an intron, disrupting *Kptn* gene function at the mRNA level by interfering with transcription downstream of the cassette site^39,107,108^.

Housing and breeding of mice and experimental procedures were carried out under the authority of UK Home Office project and personal licenses. All procedures were carried out in accordance with the Animal Welfare and Ethical Review Body of the Wellcome Sanger Institute and the UK Home Office Animals (Scientific Procedures) Act of 1986, under UK Home Office PPL 80/2472 and PPL P6320B89B. Mice were housed in specific pathogen-free mouse facilities with 12 h light/dark cycle. Food and water were available *ad libitum*, except where indicated otherwise.

### Mouse behavioural paradigms

In all assays, the mice were tracked using over-head infrared video cameras and a user-independent automated video tracking software Ethovision XT 8.5 (Noldus Information Technology, The Netherlands). All experiments were carried out on mice aged 10-22 weeks. The mice were handled for 2-3 days before the onset of testing. All mice were habituated to the behavioural room in their home cages for ≥ 30 minutes under the same light condition as the test. Male mice were tested unless stated otherwise.

### Open field assay

To test for locomotor capabilities, open field test was run as previously described^40^. In all open field trials, mice were placed in the left corner of the open field arena. Two animals were tested in parallel in two separate open fields (74 cm x 74 cm), for a total observation time of 10 minutes. A centre zone was designated with equidistant borders to the open field walls (8 cm). The arenas were cleaned thoroughly with ethanol free wipes to remove odour traces. A period of movement was defined when the mouse reached a velocity of 2 cm/s over two frames; a period of non-movement was defined when velocity was lower than 1.75 cm/s over two frames. Significant differences in distance moved and time spent moving were assessed using a Student’s t-test (α=0.05).

### Light/dark box assay

Light/dark box assay was used to test for anxiety-related behaviours, as previously described^41,42^. The assay was conducted in a darkened room. After 30 minutes of habituation to the room in their home cages, mice were separated into new cages (with clean bedding, food pellets, cardboard play tunnels, and some used bedding), with two mice per cage. The light/dark box was placed into an open field arena, positioned in the same location in the open field arena in all experiments. The light zone had two spotlights facing down directly into the compartment, illuminating the whole of the ‘light’ compartment evenly (300-400 Lux). The dark zone was sheltered from the light by an opaque lid and a door at the opening between the two compartments. The mouse was placed in the centre of the dark compartment. The door was released at the start of the trial. Trial duration was 10 minutes, and two mice were tested in parallel. The time spent in each of the compartments, as well as the frequency of the transition between zones, was recorded. Significant differences between genotypes was identified using a Student’s t-test (α=0.05). Mice that do not transition into the light zone at all during the duration of the assay are excluded due to the possibility that the apparent preference is due to a lack of exploration. One *Kptn*^−/−^ mutant was excluded from the final analysis because it remained the dark zone the whole time.

### Social recognition (SoR)

To test olfactory-mediated memory, the SoR assay was used based on previously published protocols^45,109^. Test female animals were habituated to the room (and red light) for half an hour in their home cages and were then habituated to the arenas for 10 minutes before the start of the testing. Stimuli were matched to test animals according to their weight, size, and gender. The familiar-unfamiliar stimuli pair were from different breeding colony, but of the same genetic background (129P2/OlaHsdWtsi + C57BL/6J), and counterbalanced for novelty. Mice were exposed to the same stimulus animals for four consecutive 1 min trials, with a 10 min inter-trial-interval. In trial 5 (1 min), a novel C57BL/6NTac mouse was used. For each trial, the same stimulus was used for 3 test animals to reduce the number of animals needing to be anaesthetised.

On Day 2, after a 10 min habituation trial, mice were tested in a 2 min discrimination trial in which they were simultaneously exposed to two stimulus animals, a familiar mouse previously encountered on Day 1 (Trial 1 to 4) and a novel unfamiliar mouse. The time spent closely investigating the stimulus animals was recorded. Significant differences between genotypes in investigation times were identified using two-way ANOVA with repeated measures for the trial/stimulus mouse (α=0.05). Stimulus animals were subject to non-terminal anaesthesia with ketamine/xylazine (intraperitoneal injection 100mg/10mg per kg of body weight) and were recovered with atipamezole (subcutaneous injection 1mg/kg or 0.025mg) in a home cage with a warm water bottle at the end of the experiment. The preference index is defined as the fraction of time spend investigating the unfamiliar mouse out of the total investigation time of both mice. The preference indices were compared using an unpaired, two-tailed students t-test (α=0.05). Mice were excluded if their overall investigation time was <10 seconds on trial 1 or if they did not investigate one of the stimuli during Day 2. One wild type and four mutants were excluded due to the low investigation on trial 1 on Day 1 (final: *Kptn^+/+^* n=11, *Kptn^−/−^* n=8).

### Pairwise discrimination

Pairwise visual discrimination (PD) assay was run using an automated touchscreen technology^110–112^. The protocol used was adapted from previous studies and the Campden instruction manual for PD task (for mouse touch screen systems and ABET II)^51,111,112^. Because the assay is dependent on appetitive reward, the mice were food restricted to achieve a gradual reduction of 10-15% from their initial body weight, and this weight was maintained throughout training and testing. The weight of the mice was measured daily, and the food was adjusted accordingly. All the mice were progressed through the assay on an individual basis. Mice were tested in the same testing box each time throughout all the sessions. Once each mouse reached the criteria in the PD task phase they were no longer tested. In order to represent the percentage of correct trials over sessions in PD task, for those mice that completed the PD task earlier, the criteria mean (correct trial percentage over the last two days) was calculated, and this number was included into the overall mean for the subsequent sessions. Therefore, when the percentage of trials was plotted against sessions, each session had an equal number of mice, even though some mice finished the assay faster than others. The actual number of days taken to reach criteria was plotted separately (Student’s two-tailed t-test, α=0.05). There was a predefined 25 days cut-off point, after which if mice did not complete the criteria, they were excluded.

### Barnes maze

The Barnes maze assay was run as described previously^46–48^. External spatial cues were placed around the walls of the room, to aid navigation. The mice were tested in a darkened room with overhead lights facing the Barnes maze tables (120 cm diameter with 20 x 5cm diameter holes), under maximum illumination. Three days before the first session, their cages were cleaned and mice were familiarised with the goal box, by placing one in the centre of the clean home cage. Mice were placed into the box and allowed to climb out of it. If the mouse did not leave the box after one minute, it was gently encouraged back out into the cage. The mice were tested sequentially and in the same order on every day, with two mice tested in parallel on two tables. Fifteen minutes before testing, mice were singly housed in new cages with bedding, food, a cardboard play tunnel, and a handful of old bedding from the home cage of the mouse. Mice were trained to find the goal box for two days (Training 1, four trials each day) and tested on Day 3 with a 24 h Probe trial. Animals were then retrained to a new location for 10 trials (Training 2); two trials on Day 3, four trials on Day 4 and another four trials on Day 5. On Day 8, mice were tested again with a 72 h Probe trial.

The time it took the mouse to find the goal box (primary latency), was recorded for each training trial. For the Probe trials, the time spent in the areas around each of the holes (10cm diameter area) and the average distance away from each of the holes was recorded. The time around each hole was calculated as a percentage of the total time spent around the holes (i.e. the sum of all the intervals in proximity to the holes). Significant differences between genotypes were identified using two-way ANOVA with genotype and hole/training day as factors (α=0.05).

### Transcardial perfusions

Mice were terminally anaesthetised by intraperitoneal injection of 0.1mL pentobarbitone sodium. After the mice were no longer responsive, 20 mL of cold PBS was injected into the left ventricle of the exposed heart at low speed, followed by 20 ml of 4% PFA, either by hand (20-50 ml syringe) or using a perfusion pump. After the transcardial perfusion, skulls and brain segments were collected.

### MRI (Magnetic Resonance Imaging)

For MRI analysis, skulls were collected after transcardial perfusion, and stored in formalin solution at 4C. MRI imaging and analysis were performed, using previously published methodology^57^. Brains were scanned using a Bruker PharmaScan 47/16 system at 4.7T with a manufacturer-provided birdcage transmit-receive coil. The images were registered into the same stereotactic space and segmented into images of four tissue types: grey matter, white matter, cerebrospinal fluid and ‘other’. This was followed by voxel-based quantification of brain volume^113^. To control the type I error rate due to multiple comparisons, an adjusted p-value was used for a false discovery rate at p < 0.05.

### Microcomputed X-ray tomography

Skulls of male mice (n=5 per genotype) were collected after transcardial perfusion and stored in formalin solution at 4C until they were ready to be scanned. High-resolution images were acquired using the 1172 Bruker microCT scanner. The scanning parameters were set to: 2000 x 1332 object position, pixel size 9.89, Medium camera size. Prior to acquisition, we ensured the average intensity value (av=xx) was in the acceptable range of 40-60% (no more than 70%) and the ‘min’ value was ≥10%. DICOM files generated by the Bruker software were loaded into 3DimViewer (V3.0, http://www.3dim-laboratory.cz) for surface model creation using default thresholding (1482/7000) and saved as an .STL file. The STL files were loaded individually into Meshlab software^114^ (V2016.12, www.meshlab.net), and individual landmark locations^54^ were recorded (Supplementary Fig. 3). Inter-landmark distances were calculated in excel. Significant differences in mean linear distances were identified using a Student’s t-test (α=0.05). Visualisation, 3d rendering and resectioning was performed in Meshmixer V3.5.474 (Autodesk, Inc., www.meshmixer.com).

### Histomorphometric analysis

For the histomorphological analysis, adult mice aged 16 weeks (n=8 per genotype and sex) were transcardially perfused, the brain was removed from the skull and post-fixed in formalin for 48 hours, after which it was kept in PBS. For the P0 brains, male mice were culled at birth, brains were removed from skull and post-fixed for 12 days before being transferred into PBS. The morphometric analyses were carried out using a previously published high-throughput method^58,59^, examining the same three coronal brain regions (section 1: Bregma +0.98mm; section 2: Bregma −1.34mm; and section 3: Bregma −5.80mm) in all mice. Mice were separately analysed according to their sex. Sections were scanned using Slide scanner (Hamamatsu, NanoZoomer 2.0HT, C9600 series) and accessories (racks and NanoZoomer digital pathology, version 2.5.64 software) at 20x magnification and analysed using ImageJ (78 measurements for P20 and 16-week animals, 53 measurements for P0 animals, as listed in Supplementary Table 2).

### Immunohistochemistry

Adult mice were euthanized by CO2 asphyxiation followed by thoracotomy. All animals were perfused with ice-cold saline and 4% PFA. Brains were dissected out and post-fixed in 4% PFA for 24 h. After fixation, brains were embedded in paraffin wax, sliced on a microtome at 8 μm, mounted on slides, and hot air dried overnight. Following drying, brains were deparaffinized, subjected to antigen retrieval via citrate buffer, and washed in distilled water. Washed slides were blocked for 1 h at room temperature (RT) in blocking solution containing 0.3% tween PBS with 5% normal goat serum and incubated in primary antibodies overnight in blocking solution. Primary antibodies used were anti-phospho-RPS6 (ser240/244; Cell Signaling Technology; 1:1000; mouse anti-rabbit polyclonal) and RPS6 (NSJ Bioreagents; 1:100, mouse anti-rabbit monoclonal). After incubation with primary antibodies, slides were washed in PBS and placed in blocking solution with secondary antibodies for 2 h at RT. Secondary antibodies were conjugated with Alexa 594 (Invitrogen; 1:1000; goat anti-rabbit) and 647 (Invitrogen; 1:1000; goat anti-rabbit) fluorophores, respectively. After two hours, slides were washed in PBS and distilled water and finally coverslipped with Vectashield media containing DAPI. Slides were then imaged on a Nikon spinning disk confocal microscope and digital micrographs were created in Imaris software (Oxford Instruments).

### Tissue lysis and immunoblotting

For protein extracts used in immunoblotting, PBS perfused tissue was collected as described above, snap frozen in liquid nitrogen, and stored in −80C until lysis. Tissue was lysed in 1x lysis solution (protease + phosphatase inhibitor cocktail in TPER) in the Qiagen TissueLyser LT with sterile steel beads and operated at 30Hz for 2 minutes. Samples were spun at 4C for 15 minutes at 15000 RPM and the supernatant was kept on ice. Bio-Rad DC protein assay kit was used for protein quantifications, following the manufacturer′s protocol. Samples were boiled for 5 minutes and then subjected to XCell4 SureLock Midi-Cell electrophoresis. 20uL of 30ug of protein was run per well. Membranes were blocked for an hour in 5% BSA (in PBS with 0.1% Tween). Primary antibodies were incubated overnight at 4C, followed by 1 h incubation in horseradish peroxidase–conjugated secondary antibody (anti-rabbit or anti-mouse, #7074 and #7076, Cell Signaling Technology) diluted in PBS-T 5% milk at room temperature. Images were acquired with a digital imaging system (ImageQuant LAS 4000, GE Healthcare) and analysed using ImageJ software. Primary antibodies: anti-RPS6 (clone 54D2, #2317, Cell Signaling Technology), Ser240/244-specific anti-phospho-RPS6 (#2215, Cell Signaling Technology), and anti-GAPDH (ab9485, Abcam).

### Rapamycin treatment

We freshly dissolved stock rapamycin (dissolved in ethanol 20mg/mL (LC Laboratories)) in vehicle solution (5% Tween, 5% PEG, ethanol, in distilled water) before use. We administered rapamycin intraperitoneally once daily at a dose of 5 mg/kg for 3 days before and on the day of collecting tissue for immunoblotting.

### Mutagenesis and differentiation of KPTN iPSC models

iPSC experiments were all performed in an isogenic background: Kolf2C1_WT (HPSI0114i-kolf_2), feeder-free hiPSC, male, derived from skin tissue using Cytotune 1 reprogramming^76^. In order to create knockout iPSC lines, a guide RNA (gRNA) was selected to target coding exon 8 of the human KPTN gene (GRCh38:19:47479882-47479904). The synthetic gRNA along with Cas9 protein was delivered into wild-type Kolf2C1 cells^76^ as a pre-complexed ribonucleoprotein via electroporation. The addition of a short single-stranded oligodeoxynucleotide of non-complementary sequence was also added to improve delivery. After a period of recovery, the cells were subcloned and submitted for genotyping by Miseq sequencing. Clones were screened for the presence of frameshift-causing indels and then expanded for banking. Clones identified as unedited in the genotyping analysis were selected as unedited controls, to control for electroporation, selection, and passaging effects. After banking, and subsequent to all differentiation experiments, all cell lines were sent for a second round of confirmatory genotyping. This work was performed by the Gene Editing facility at the Wellcome Sanger Institute.

### Differentiation of iPSCs to cortical neural precursor cells

*KPTN* mutant heterozygous, homozygous edited and wild-type control IPSC lines were cultured in feeder-free conditions as per published protocols^77^. Dual-SMAD inhibition (SB431542, #ab120163, Abcam and LDN193189, #2092-5, Cambridge Bioscience) was used in a 10-day neural induction in the presence of XAV939 (#3748, Tocris) to promote regional forebrain identity^77^. After passaging at day 10, cells were replated for immunocytochemical quality control, and then banked at day 14 for RNA extraction and RNA-Seq analysis. All experiments were confirmed to have high PAX6/GFAP positivity (>90% of cells at day 14). All wildtype, heterozygous, and homozygous clones (n=2,2,1 respectively) were differentiated in duplicate, providing 2-4 separate biological replicates per genotype (n=4,4,2 respectively).

### iPSC RNA extraction and RNA sequencing

RNA extraction and sequencing of iPSC derived materials was performed using manufacturer’s protocols for the RNeasy QIAcube kit on a QIAcube automated system (Qiagen). RNA sequencing libraries were prepared using established protocols: library construction (poly(A) pulldown, fragmentation, 1st and 2nd strand synthesis, end prep and ligation) was performed using the NEB Ultra II RNA custom kit (New England Biolabs) on an Agilent Bravo automated system. Indexed multiplexed sequencing was performed on the Novaseq 6000 system (S4 flow cell, Xp workflow; Illumina), collecting approximately 30 million paired-end reads per sample with 100 base read length. The sequencing data were de-multiplexed into separate CRAM files for each library in a lane. Adapters that had been hard-clipped prior to alignment were reinserted as soft-clipped post alignment, and duplicated fragments were marked in the CRAM files. The data pre-processing, including sequences QC, and STAR alignments were made with a custom Nextflow pipeline, which is publicly available at https://github.com/wtsi-hgi/nextflow-pipelines/blob/rna_seq_5607/pipelines/rna_seq.nf, including the specific aligner parameters. We assessed the sequence data quality using FastQC v0.11.8. Reads were aligned to the GRCh38 human reference genome (Ensembl GTF annotation v91). We used STAR^115^ version STAR_2.6.1d with the --twopassMode Basic parameter. The STAR index was built against GRCh38 Ensembl GTF v91 using the option -sjdbOverhang 75. We then used featureCounts version 1.6.4^116^ to obtain a readcount matrix. Genes with no count or only a single count across all samples were filtered out. The counts were normalised using DESEQ2’s median of ratios method^117^. Differential gene expression was analysed using the DESEQ2 package^117^ with SVA correction^118^. An adjusted p-value threshold of 0.05 was selected to identify significant differences between wild-type and mutant samples.

### Mouse RNA extraction and RNA sequencing

For wild-type and homozygous *Kptn* mouse samples, we used n=5-6 per genotype and tissue as follows (wildtype/mutant): E18 brain, n=6/6; P21 hippocampus, n=5/6; P21 cortex, n=5/6; adult cerebellum, n=5/6; adult hippocampus, n=6/6; adult cortex, n=5/5. Fresh frozen mouse tissues were homogenised in buffer RLT plus (Qiagen) with β-mercaptoethanol (Sigma, M3148; 10µl/ml) using Qiagen TissueLyser LT, with sterile RNAse-ZAP treated steel beads and operated at 50Hz for 2 minutes. Samples were pre-treated on gDNA eliminator columns and then extracted on RNeasy Plus columns as per manufacturer’s protocol (Qiagen), and were immediately snap frozen on dry ice and stored at −80C. An aliquot of each sample was quantified using 2100 Bioanalyzer (Agilent Technologies). Library preparation was performed by Wellcome Sanger Institute DNA Pipelines, as described above for iPSCs. Samples were sequenced using 75 bp paired-end sequencing reads (reverse stranded) on a Illumina-HTP HiSeq 4000 system. The sequencing data were de-multiplexed into separate CRAM files for each library in a lane. Adapters that had been hard-clipped prior to alignment were reinserted as soft-clipped post alignment, and duplicated fragments were marked in the CRAM files. The data pre-processing, including sequences QC, and STAR alignments were made with a Nextflow pipeline, which is publicly available at https://github.com/wtsi-hgi/nextflow-pipelines/blob/rna_seq_mouse/pipelines/rna_seq.nf, including the specific aligner parameters. We assessed the sequence data quality using FastQC v0.11.8. Reads were aligned to the GRCm38 mouse reference genome (Mus_musculus.GRCm38.dna.primary_assembly.fa, Ensembl GTF annotation v99). We used STAR version 2.7.3a^115^ with the --twopassMode Basic parameter. The STAR index was built against Mus_musculus GRCm38 v99 Ensembl GTF using the option -sjdbOverhang 75. We then used featureCounts version 2.0.0^116^ to obtain a readcount matrix. The count data was used as input for differential gene expression analysis using DESeq2 package^117^ with SVA correction^118^. The default DESeq2 cut-off of BH-adjusted p-value<0.1 was used for the mouse RNA-Seq analyses.

For the identification of functionally enriched terms in the differentially expressed genes, Gene ontology (GO) and KEGG pathway enrichment analyses was performed using the grpofiler online suite^72^ (http://biit.cs.ut.ee/gprofiler/index.cgi). A threshold of 5% FDR and an enrichment significance threshold of p<0.05 (hypergeometric test with Benjamini-Hochberg False Discovery rate correction for multiple testing) was used. In all analyses, a background comprised of only the expressed genes in the tissue studied (genes where the adjusted p-value yielded a numerical value, different to NA).

### Clinical and genetic studies

Affected individuals were identified by their clinician, using GeneMatcher^119^ and through the published medical literature. Research was performed with informed consent from the study participants or their legal guardians, according to institutional and international guidelines for studies with human participants and materials. Institutional study approvals include: Akron Children’s Hospital (IRB 986876-3), University of Arizona (IRB 10-0050-010), University of Exeter Medical School. Affected individuals and family members were investigated according to routine clinical standards for the diagnosis of developmental delay/intellectual disability and neurological disease. Genetic studies were performed as part of clinical and/or research investigations dependent on clinical presentation and family history. Detailed phenotype and genotype information was obtained by the clinical care provider using a targeted questionnaire to identify specific clinical features including occipitofrontal circumference (OFC). All OFC z-scores were recalculated using measurements and known age at assessment using the British 1990 growth reference^120^ to ensure all values were comparable. Z-score conversions around birth are gestation-adjusted where needed.

### Psychometric testing of *KPTN*-related disorder probands

Six Amish individuals between the ages of 11 and 29 with KRD (3 female, 3 male) were psychometrically assessed using the Wechsler Intelligence Scale for Children 4th Edition (WISC-IV) (6-16 years) or the Wechsler Adult Intelligence Scale 4^th^ Edition (WAIS-IV) (over 16 years), which assesses cognitive performance in four domains including verbal comprehension (VCI), perceptual reasoning (PRI), processing speed (PSI) and working memory (WMI) indices, and can be combined to generate a full-scale intelligence quotient (FSIQ) score. Immediate memory recall was assessed using the story memory test and list learning test from the developmental NEuroPSYchological Assessment-2^nd^ Edition (NEPSY-II) (5-16 years) or the Repeatable Battery for Assessment of Neurological Status-2^nd^ Edition (RBANS-Update) (over 12 years). All researchers collecting assessment data were trained and supervised by a Clinical Psychologist. The WISC-IV and WAIS-IV tests have normalised reference scores, with mean of 100 and an SD of 15, although to allow comparison standardised scores were converted to Z-scores. The NEPSY-II and RBANS-Update tests generate scaled scores, with mean of 10 and an SD of 3, which were also converted to Z-scores for direct comparison. As none of these tests have been validated within the Amish population and due to possible cultural differences that could affect test scores, this necessitated data collection from a reference control group. Six age-matched Amish control individuals (6 female) who were unaffected siblings or relatives of the six affected individuals, underwent the same psychometric tests, to serve as controls for this analysis.

### CMAP Query

To identify anti-correlated transcriptional profiles in the CMAP database^121^, we selected the 150 most strongly up- and down-regulated genes from our differential expression analysis of KPTN^−/−^ NPCs, and queried the CMAP via their online tool (https://clue.io/query) using the L1000 TOUCHSTONE (V1.1.1.2) gene expression dataset.

## Supplementary materials

Supplementary data legends.pdf

SUPP FIGURE2 pedigree.pdf

SUPP FIGURE1 EXPRESSION.pdf

SUPP FIGURE3 uCT Landmarks.pdf

SUPP FIGURE4 ANATOMY.pdf

SUPP FIGURE5 OFC Proband 14.pdf

SUPP FIGURE6 CMAP.pdf

SUPP FIGURE7 SEIZURE.pdf

SUPP FILE1 MOUSE PATHWAY ANALYSIS.xlsx

SUPP TABLE1 uCT DATA.pdf

SUPP TABLE2 ANATOMY.xlsx

SUPP TABLE3 KRD PROBANDS.xlsx

SUPP TABLE4 RNASEQ.xlsx

SUPP TABLE5 KPTN cell types analysis.pdf

SUPP TABLE6 DD GENE PHENOTYPES.pdf

## Author contributions

D.W.L., A.C., E.L.B., M.O.L., and S.S.G. conceived of the original project; M.O.L., G.S.A., D.W.L., M.E.H., and S.S.G. designed mouse experiments; D.W.L., M.E.H., and S.S.G. supervised the project; E.L.Cambridge, L.T., and M.S. performed initial mouse phenotyping; C.J.L. supervised initial mouse phenotyping; M.O.L, D.W.L., G.S.A., E.L.Coomber, M.S., C.R., and S.S.G carried out mouse behavioural phenotyping; L.E.R., E.L.B., and A.C. supervised clinical data collection, and psychometric testing; J.T. and L.J. performed psychometric testing and analysis; C.B., I.E., J.C.H., S.G.K., E.N., A.Y.S., E.M.S., H.S., J.S., T.T., C.V.R.A., P.V., M.W., and O.W. provided clinical data; L.E.R., and E.L.B. collated clinical data; M.O.L., G.S.A., and H.I. performed sample preparation, western blotting, and qPCR; S.C.C. performed mouse histological processing and analysis; B.Y. supervised mouse histological brain analysis; M.O.L. carried out μCT scans; S.S.G. analysed μCT data; P.H.I.II performed mouse immunochemistry; P.B.C. supervised mouse immunochemistry, and contributed to manuscript preparation; S.J.S. carried out MRI scanning and analysis; A.D., A.H., and M.P. supervised iPSC work; M.H.L.A, A.N., and E.R. carried out iPSC differentiations; M.O.L., G.S.A., M.S., and S.S.G analysed mouse behavioural data; G.S.A., O.A., and S.S.G. analysed RNA-Seq results; M.O.L., G.S.A., L.E.R., E.L.B., and S.S.G. wrote the manuscript, and assembled figures.

## References

1. Azevedo, F. A. C. et al. Equal numbers of neuronal and nonneuronal cells make the human brain an isometrically scaled-up primate brain. J. Comp. Neurol. 513, 532–541 (2009).

2. Walsh, C. A. & Engle, E. C. Allelic Diversity in Human Developmental Neurogenetics: Insights into Biology and Disease. Neuron 68, 245–253 (2010).

3. Zoghbi, H. Y. Postnatal neurodevelopmental disorders: meeting at the synapse? Science (80-.). 302, 826–30 (2003).

4. Rubenstein, J. L. R. Annual Research Review: Development of the cerebral cortex: implications for neurodevelopmental disorders. J. Child Psychol. Psychiatry 52, 339– 355 (2011).

5. Wright, C. F. et al. Genetic diagnosis of developmental disorders in the DDD study: A scalable analysis of genome-wide research data. Lancet 385, 1305–1314 (2015).

6. Firth, H. V et al. DECIPHER: Database of Chromosomal Imbalance and Phenotype in Humans Using Ensembl Resources. Am. J. Hum. Genet. 84, 524–33 (2009).

7. Clark, R. D. & Curry, C. J. Macrocephaly and Megalencephaly. in Genetic Consultations in the Newborn (eds. Clark, R. D. & Curry, C. J.) (Oxford University Press, 2019). doi:10.1093/med/9780199990993.003.0014.

8. Winden, K., Yuskaitis, C. & Poduri, A. Megalencephaly and Macrocephaly. Semin. Neurol. 35, (2015).

9. Keppler-Noreuil, K. M., Parker, V. E. R., Darling, T. N. & Martinez-Agosto, J. A. Somatic overgrowth disorders of the PI3K/AKT/mTOR pathway & therapeutic strategies. Am. J. Med. Genet. Part C Semin. Med. Genet. 172, 402–421 (2016).

10. Lipton, J. O. & Sahin, M. The neurology of mTOR. Neuron 84, 275–91 (2014).

11. Saxton, R. A. & Sabatini, D. M. mTOR Signaling in Growth, Metabolism, and Disease. Cell 168, 960–976 (2017).

12. Tee, A. R., Sampson, J. R., Pal, D. K. & Bateman, J. M. The role of mTOR signalling in neurogenesis, insights from tuberous sclerosis complex. Seminars in Cell and Developmental Biology vol. 52 12–20 (2016).

13. Crino, P. B. mTOR: A pathogenic signaling pathway in developmental brain malformations. Trends Mol. Med. 17, 734–42 (2011).

14. Hoeffer, C. A. & Klann, E. mTOR signaling: At the crossroads of plasticity, memory and disease. Trends in Neurosciences vol. 33 67–75 (2010).

15. Frankel, W. N., Yang, Y., Mahaffey, C. L., Beyer, B. J. & O’Brien, T. P. Szt2, a novel gene for seizure threshold in mice. Genes. Brain. Behav. 8, 568–76 (2009).

16. Crino, P. B. MTOR: A pathogenic signaling pathway in developmental brain malformations. Trends Mol. Med. 17, 734–742 (2011).

17. Basel-Vanagaite, L. et al. Biallelic SZT2 mutations cause infantile encephalopathy with epilepsy and dysmorphic corpus callosum. Am. J. Hum. Genet. 93, 524–9 (2013).

18. Negishi, Y. et al. A combination of genetic and biochemical analyses for the diagnosis of PI3K-AKT-mTOR pathway-associated megalencephaly. BMC Med. Genet. 18, (2017).

19. Nakamura, Y. et al. Constitutive activation of mTORC1 signaling induced by biallelic loss-of-function mutations in SZT2 underlies a discernible neurodevelopmental disease. PLoS One 14, e0221482 (2019).

20. Dang, L. T. et al. Multimodal Analysis of STRADA Function in Brain Development. Front. Cell. Neurosci. 14, (2020).

21. Pollen, A. A. et al. Establishing Cerebral Organoids as Models of Human-Specific Brain Evolution. Cell 176, 743–756.e17 (2019).

22. Hartman, N. W. et al. mTORC1 targets the translational repressor 4E-BP2, but not S6 kinase 1/2, to regulate neural stem cell self-renewal in vivo. Cell Rep. 5, 433–44 (2013).

23. Ka, M., Condorelli, G., Woodgett, J. R. & Kim, W. Y. mTOR regulates brain morphogenesis by mediating GSK3 signaling. Development 141, 4076–4086 (2014).

24. Meng, D., Frank, A. R. & Jewell, J. L. mTOR signaling in stem and progenitor cells. Development 145, (2018).

25. Lee, J. H. et al. De novo somatic mutations in components of the PI3K-AKT3-mTOR pathway cause hemimegalencephaly. Nat. Genet. 44, 941–5 (2012).

26. D’Gama, A. M. et al. Mammalian target of rapamycin pathway mutations cause hemimegalencephaly and focal cortical dysplasia. Ann. Neurol. 77, 720–5 (2015).

27. Gordo, G. et al. mTOR mutations in Smith-Kingsmore syndrome: Four additional patients and a review. Clin. Genet. 93, 762–775 (2018).

28. Baple, E. L. et al. Mutations in KPTN cause macrocephaly, neurodevelopmental delay, and seizures. Am. J. Hum. Genet. 94, 87–94 (2014).

29. Pajusalu, S., Reimand, T. & Õunap, K. Novel homozygous mutation in KPTN gene causing a familial intellectual disability-macrocephaly syndrome. Am. J. Med. Genet. A 167A, 1913–5 (2015).

30. Lucena, P. H. et al. *kptn* gene homozygous variant-related syndrome in the northeast of Brazil: A case report. Am. J. Med. Genet. Part A 182, 762–767 (2020).

31. Pacio Miguez, M. et al. Pathogenic variants in KPTN, a rare cause of macrocephaly and intellectual disability. Am. J. Med. Genet. A 182, 2222–2225 (2020).

32. Thiffault, I. et al. Pathogenic variants in KPTN gene identified by clinical whole-genome sequencing. Cold Spring Harb. Mol. Case Stud. 6, a003970 (2020).

33. Wolfson, R. L. et al. KICSTOR recruits GATOR1 to the lysosome and is necessary for nutrients to regulate mTORC1. Nature 543, 438–442 (2017).

34. Peng, M., Yin, N. & Li, M. O. SZT2 dictates GATOR control of mTORC1 signalling. Nature 543, 433–437 (2017).

35. Barton, A. R., Hujoel, M. L. A., Mukamel, R. E., Sherman, M. A. & Loh, P.-R. A spectrum of recessiveness among Mendelian disease variants in UK Biobank. Am. J. Hum. Genet. (2022) doi:10.1016/j.ajhg.2022.05.008.

36. Kingdom, R. et al. Rare genetic variants in genes and loci linked to dominant monogenic developmental disorders cause milder related phenotypes in the general population. Am. J. Hum. Genet. (2022) doi:10.1016/j.ajhg.2022.05.011.

37. Huang, Y. et al. Rare genetic variants impact muscle strength. medRxiv 2022.06.22.22276715 (2022) doi:10.1101/2022.06.22.22276715.

38. Chen, C.-Y. et al. The impact of rare protein coding genetic variation on adult cognitive function. medRxiv 2022.06.24.22276728 (2022) doi:10.1101/2022.06.24.22276728.

39. White, J. K. et al. Genome-wide generation and systematic phenotyping of knockout mice reveals new roles for many genes. Cell 154, 452–64 (2013).

40. Seibenhener, M. L. & Wooten, M. C. Use of the Open Field Maze to measure locomotor and anxiety-like behavior in mice. J. Vis. Exp. e52434 (2015) doi:10.3791/52434.

41. Crawley, J. & Goodwin, F. K. Preliminary report of a simple animal behavior model for the anxiolytic effects of benzodiazepines. Pharmacol. Biochem. Behav. 13, 167–70 (1980).

42. Hascoët, M. & Bourin, M. The mouse light-dark box test. Neuromethods vol. 42 197– 223 (2009).

43. Ferguson, J. N., Aldag, J. M., Insel, T. R. & Young, L. J. Oxytocin in the medial amygdala is essential for social recognition in the mouse. J. Neurosci. 21, 8278–85 (2001).

44. Winslow, J. T. & Insel, T. R. Neuroendocrine basis of social recognition. Current Opinion in Neurobiology vol. 14 248–253 (2004).

45. Sánchez-Andrade, G., James, B. M. & Kendrick, K. M. Neural Encoding of Olfactory Recognition Memory. J. Reprod. Dev. 51, (2005).

46. Koopmans, G., Blokland, A., van Nieuwenhuijzen, P. & Prickaerts, J. Assessment of spatial learning abilities of mice in a new circular maze. Physiol. Behav. 79, 683–93 (2003).

47. Harrison, F. E., Reiserer, R. S., Tomarken, A. J. & McDonald, M. P. Spatial and nonspatial escape strategies in the Barnes maze. Learn. Mem. 13, 809–19 (2006).

48. Harrison, F. E., Hosseini, A. H. & McDonald, M. P. Endogenous anxiety and stress responses in water maze and Barnes maze spatial memory tasks. Behav. Brain Res. 198, 247–251 (2009).

49. Chudasama, Y. & Robbins, T. W. Dissociable contributions of the orbitofrontal and infralimbic cortex to pavlovian autoshaping and discrimination reversal learning: further evidence for the functional heterogeneity of the rodent frontal cortex. J. Neurosci. 23, 8771–80 (2003).

50. Bussey, T. J. J. et al. New translational assays for preclinical modelling of cognition in schizophrenia: The touchscreen testing method for mice and rats. Neuropharmacology 62, 1191–1203 (2012).

51. Brigman, J. L. et al. GluN2B in corticostriatal circuits governs choice learning and choice shifting. Nat. Neurosci. 16, 1101–10 (2013).

52. Izquierdo, A. et al. Basolateral amygdala lesions facilitate reward choices after negative feedback in rats. J. Neurosci. 33, 4105–9 (2013).

53. Yarkoni, T., Speer, N. K. & Zacks, J. M. Neural substrates of narrative comprehension and memory. Neuroimage 41, 1408–1425 (2008).

54. de Carlos, F. et al. 3D-μCT Cephalometric Measurements in Mice. in Computed Tomography - Special Applications (InTech, 2011). doi:10.5772/24234.

55. Xu, H. et al. Dental and craniofacial defects in the Crtap−/− mouse model of osteogenesis imperfecta type VII. Dev. Dyn. 249, 884–897 (2020).

56. Baddam, P. et al. Neural crest-specific loss of Bmp7 leads to midfacial hypoplasia, nasal airway obstruction and disordered breathing, modeling obstructive sleep apnea. Dis. Model. Mech. 14, (2021).

57. Sawiak, S. J., Wood, N. I., Williams, G. B., Morton, A. J. & Carpenter, T. A. Voxel-based morphometry with templates and validation in a mouse model of Huntington’s disease. Magn. Reson. Imaging 31, 1522–1531 (2013).

58. Mikhaleva, A., Kannan, M., Wagner, C. & Yalcin, B. Histomorphological Phenotyping of the Adult Mouse Brain. Curr. Protoc. Mouse Biol. 6, 307–332 (2016).

59. Collins, S. C. et al. Large-scale neuroanatomical study uncovers 198 gene associations in mouse brain morphogenesis. Nat. Commun. 10, 3465 (2019).

60. Kwon, C.-H. et al. Pten regulates neuronal soma size: a mouse model of Lhermitte-Duclos disease. Nat. Genet. 29, 404–411 (2001).

61. Klintsova, A., Hamilton, G. & Boschen, K. Long-Term Consequences of Developmental Alcohol Exposure on Brain Structure and Function: Therapeutic Benefits of Physical Activity. Brain Sci. 3, 1–38 (2012).

62. Semple, B. D., Blomgren, K., Gimlin, K., Ferriero, D. M. & Noble-Haeusslein, L. J. Brain development in rodents and humans: Identifying benchmarks of maturation and vulnerability to injury across species. Progress in Neurobiology vols 106-107 1–16 (2013).

63. Crino, P. B. mTOR signaling in epilepsy: Insights from malformations of cortical development. Cold Spring Harb. Perspect. Med. 5, a022442 (2015).

64. Tatton-Brown, K. & Weksberg, R. Molecular Mechanisms of Childhood Overgrowth. Am. J. Med. Genet. Part C Semin. Med. Genet. 163, 71–75 (2013).

65. Klein, S. D. et al. Mutations in the sonic hedgehog pathway cause macrocephaly-associated conditions due to crosstalk to the PI3K/AKT/mTOR pathway. Am. J. Med. Genet. Part A 179, 2517–2531 (2019).

66. Ruvinsky, I. & Meyuhas, O. Ribosomal protein S6 phosphorylation: from protein synthesis to cell size. Trends in Biochemical Sciences vol. 31 342–348 (2006).

67. Biever, A., Valjent, E. & Puighermanal, E. Ribosomal protein S6 phosphorylation in the nervous system: From regulation to function. Front. Mol. Neurosci. 8, 75 (2015).

68. Ballou, L. M. & Lin, R. Z. Rapamycin and mTOR kinase inhibitors. J. Chem. Biol. 1, 27–36 (2008).

69. Spilman, P. et al. Inhibition of mTOR by rapamycin abolishes cognitive deficits and reduces amyloid-beta levels in a mouse model of Alzheimer’s disease. PLoS One 5, e9979 (2010).

70. Way, S. W. et al. The differential effects of prenatal and/or postnatal rapamycin on neurodevelopmental defects and cognition in a neuroglial mouse model of tuberous sclerosis complex. Hum. Mol. Genet. 21, 3226–3236 (2012).

71. Rozengurt, E., Soares, H. P. & Sinnet-Smith, J. Suppression of feedback loops mediated by pi3k/mtor induces multiple overactivation of compensatory pathways: An unintended consequence leading to drug resistance. Molecular Cancer Therapeutics vol. 13 2477–2488 (2014).

72. Raudvere, U. et al. g:Profiler: a web server for functional enrichment analysis and conversions of gene lists (2019 update). Nucleic Acids Res. 47, W191–W198 (2019).

73. Cardenas, M. E., Cutler, N. S., Lorenz, M. C., Di Como, C. J. & Heitman, J. The TOR signaling cascade regulates gene expression in response to nutrients. Genes Dev. 13, 3271–9 (1999).

74. Mayer, C. & Grummt, I. Ribosome biogenesis and cell growth: mTOR coordinates transcription by all three classes of nuclear RNA polymerases. Oncogene vol. 25 6384–6391 (2006).

75. Wang, X. & Proud, C. G. The mTOR pathway in the control of protein synthesis. Physiology vol. 21 362–369 (2006).

76. Kilpinen, H. et al. Common genetic variation drives molecular heterogeneity in human iPSCs. Nature 546, 370–375 (2017).

77. Qi, Y. et al. Combined small-molecule inhibition accelerates the derivation of functional cortical neurons from human pluripotent stem cells. Nat. Biotechnol. 35, (2017).

78. Huang, J. & Manning, B. D. The TSC1-TSC2 complex: a molecular switchboard controlling cell growth. Biochem. J. 412, 179–90 (2008).

79. Winden, K. D. et al. Biallelic mutations in TSC2 lead to abnormalities associated with cortical tubers in human ipsc-derived neurons. J. Neurosci. 39, 9294–9305 (2019).

80. Lamb, J. et al. The Connectivity Map: using gene-expression signatures to connect small molecules, genes, and disease. Science 313, 1929–35 (2006).

81. Liu, Q. et al. Characterization of Torin2, an ATP-competitive inhibitor of mTOR, ATM, and ATR. Cancer Res. 73, 2574–86 (2013).

82. Andrews, M. G., Subramanian, L. & Kriegstein, A. R. mTOR signaling regulates the morphology and migration of outer radial glia in developing human cortex. Elife 9, (2020).

83. Knobloch, M. et al. SPOT14-positive neural stem/progenitor cells in the hippocampus respond dynamically to neurogenic regulators. Stem cell reports 3, 735–42 (2014).

84. Knobloch, M. et al. Metabolic control of adult neural stem cell activity by Fasn-dependent lipogenesis. Nature 493, 226–30 (2013).

85. Shin, J. et al. Single-Cell RNA-Seq with Waterfall Reveals Molecular Cascades underlying Adult Neurogenesis. Cell Stem Cell 17, 360–72 (2015).

86. Hochgerner, H., Zeisel, A., Lönnerberg, P. & Linnarsson, S. Conserved properties of dentate gyrus neurogenesis across postnatal development revealed by single-cell RNA sequencing. Nat. Neurosci. 21, 290–299 (2018).

87. Di Bella, D. J. et al. Molecular logic of cellular diversification in the mouse cerebral cortex. Nature 595, 554–559 (2021).

88. Frankel, W. N., Yang, Y., Mahaffey, C. L., Beyer, B. J. & O’Brien, T. P. Szt2, a novel gene for seizure threshold in mice. Genes. Brain. Behav. 8, 568–76 (2009).

89. Basel-Vanagaite, L. et al. Biallelic SZT2 mutations cause infantile encephalopathy with epilepsy and dysmorphic corpus callosum. Am. J. Hum. Genet. 93, 524–9 (2013).

90. Wang, J. et al. Epilepsy-associated genes. Seizure 44, 11–20 (2017).

91. Thormann, A. et al. Flexible and scalable diagnostic filtering of genomic variants using G2P with Ensembl VEP. Nat. Commun. 10, 2373 (2019).

92. Maljevic, S. et al. Novel KCNQ3 Mutation in a Large Family with Benign Familial Neonatal Epilepsy: A Rare Cause of Neonatal Seizures. Mol. Syndromol. 7, 189–196 (2016).

93. Rogawski, M. A. KCNQ2/KCNQ3 K+ channels and the molecular pathogenesis of epilepsy: implications for therapy. Trends Neurosci. 23, 393–8 (2000).

94. Hansen, J. et al. De novo mutations in SIK1 cause a spectrum of developmental epilepsies. Am. J. Hum. Genet. 96, 682–90 (2015).

95. Pröschel, C. et al. Epilepsy-causing sequence variations in SIK1 disrupt synaptic activity response gene expression and affect neuronal morphology. Eur. J. Hum. Genet. 25, 216–221 (2017).

96. Switon, K., Kotulska, K., Janusz-Kaminska, A., Zmorzynska, J. & Jaworski, J. Molecular neurobiology of mTOR. Neuroscience 341, 112–153 (2017).

97. Kaplanis, J. et al. Evidence for 28 genetic disorders discovered by combining healthcare and research data. Nature 586, 757–762 (2020).

98. Parker, W. E. et al. Rapamycin prevents seizures after depletion of STRADA in a rare neurodevelopmental disorder. Sci. Transl. Med. 5, 182ra53 (2013).

99. French, J. A. et al. Adjunctive everolimus therapy for treatment-resistant focal-onset seizures associated with tuberous sclerosis (EXIST-3): a phase 3, randomised, double-blind, placebo-controlled study. Lancet 388, (2016).

100. Lim, J. S. et al. Brain somatic mutations in MTOR cause focal cortical dysplasia type II leading to intractable epilepsy. Nat. Med. 21, (2015).

101. Poduri, A. et al. Somatic Activation of AKT3 Causes Hemispheric Developmental Brain Malformations. Neuron 74, (2012).

102. Karczewski, K. J. et al. The mutational constraint spectrum quantified from variation in 141,456 humans. Nature 581, 434–443 (2020).

103. Reijnders, M. R. F. et al. Variation in a range of mTOR-related genes associates with intracranial volume and intellectual disability. Nat. Commun. 8, (2017).

104. Ehninger, D. et al. Reversal of learning deficits in a Tsc2+/− mouse model of tuberous sclerosis. Nat. Med. 14, 843–8 (2008).

105. Kwon, C.-H., Zhu, X., Zhang, J. & Baker, S. J. mTor is required for hypertrophy of Pten-deficient neuronal soma in vivo. Proc. Natl. Acad. Sci. U. S. A. 100, 12923–8 (2003).

106. Ehninger, D., Li, W., Fox, K., Stryker, M. P. & Silva, A. J. Reversing Neurodevelopmental Disorders in Adults. Neuron vol. 60 950–960 (Cell Press, 2008).

107. Skarnes, W. C. et al. A conditional knockout resource for the genome-wide study of mouse gene function. Nature 474, 337–342 (2011).

108. Testa, G. et al. A reliable lacZ expression reporter cassette for multipurpose, knockout-first alleles. Genesis 38, 151–8 (2004).

109. Dias, C. et al. BCL11A Haploinsufficiency Causes an Intellectual Disability Syndrome and Dysregulates Transcription. Am. J. Hum. Genet. 99, 253–274 (2016).

110. Bubser, M. et al. Selective Activation of M _4_ Muscarinic Acetylcholine Receptors Reverses MK-801-Induced Behavioral Impairments and Enhances Associative Learning in Rodents. ACS Chem. Neurosci. 5, 920–942 (2014).

111. Horner, A. E. et al. The touchscreen operant platform for testing learning and memory in rats and mice. Nat. Protoc. 8, 1961–1984 (2013).

112. Morton, A. J., Skillings, E., Bussey, T. J. & Saksida, L. M. Measuring cognitive deficits in disabled mice using an automated interactive touchscreen system. Nat. Methods 3, 767 (2006).

113. Ashburner, J. & Friston, K. J. Voxel-based morphometry--the methods. Neuroimage 11, 805–21 (2000).

114. Cignoni, P. et al. MeshLab: An open-source mesh processing tool. 6th Eurographics Ital. Chapter Conf. 2008 - Proc. 129–136 (2008).

115. Dobin, A. et al. STAR: ultrafast universal RNA-seq aligner. Bioinformatics 29, 15–21 (2013).

116. Liao, Y., Smyth, G. K. & Shi, W. featureCounts: an efficient general purpose program for assigning sequence reads to genomic features. Bioinformatics 30, 923–30 (2014).

117. Love, M. I., Huber, W. & Anders, S. Moderated estimation of fold change and dispersion for RNA-seq data with DESeq2. Genome Biol. 15, 550 (2014).

118. Leek, J. T. & Storey, J. D. Capturing heterogeneity in gene expression studies by surrogate variable analysis. PLoS Genet. 3, 1724–35 (2007).

119. Sobreira, N., Schiettecatte, F., Valle, D. & Hamosh, A. GeneMatcher: a matching tool for connecting investigators with an interest in the same gene. Hum. Mutat. 36, 928–30 (2015).

120. Cole, T. J., Freeman, J. V & Preece, M. A. British 1990 growth reference centiles for weight, height, body mass index and head circumference fitted by maximum penalized likelihood. Stat. Med. 17, 407–29 (1998).

121. Subramanian, A. et al. A Next Generation Connectivity Map: L1000 Platform and the First 1,000,000 Profiles. Cell 171, 1437–1452.e17 (2017).

