## Supplementary data legends for "Mouse and cellular models of *KPTN*-related disorder implicate mTOR signalling in cognitive and progressive overgrowth phenotypes"

### SUPPLEMENTARY FIGURE LEGENDS

**SUPPLEMENTARY FIGURE 1 *Kptn* gene expression in mouse and human loss-of-function models. a** Alignment of Exon 8 *KPTN* alleles reported in Baple *et al.* 2014 (c.714_731dup and c.776C>A), and in hIPSCs used in this study (allele 1: c717_c741del + c751_757del, allele 2: c751_757del, allele 3: c734_753del). All coordinates refer to MANE select transcript (NM_007059.4). Duplicated regions are highlighted in red, missense changes in red font, and deletions represented with dashes. **b** Normalised RNA-Seq read counts for *Kptn* in embryonic (E18 brain) and adult (Hippocampus, Cortex, Cerebellum) samples of *Kptn*$^{-/-}$ mice and *Kptn*$^{+/+}$ controls show near-compete loss of expression of the targeted gene in *Kptn* KO mice. **c** Normalised RNA-Seq read counts for *KPTN* in differentiated neural precursor cells of *Kptn*$^{+/+}$, *Kptn*$^{+/-}$, and *Kptn*$^{-/-}$ genotypes show dosage-associated loss of expression of the targeted gene.

**SUPPLEMENTARY FIGURE 2 Pedigree of an extended interrelated Amish family.** Relationship between seven individuals with KRD who underwent psychometric testing (shaded symbols). Affected individual X:2 (blue shaded symbol); psychometric testing was attempted but was not possible due to the severity of the intellectual impairment. Genotype is shown in red under each individual (X, p.(Ser259*) allele; Dup, p.(Met241_Gln246dup) allele; WT, wild-type allele).

**SUPPLEMENTARY FIGURE 3 Landmarks used for microcomputed tomography analysis.** L1-L11 landmarks were used to measure distances in adult mouse skulls of KRD models, as computed in Supplementary Table 1.

**SUPPLEMENTARY FIGURE 4 Inactivation of *Kptn* gives rise to major neuroanatomical defects. a** Neuroanatomical findings in adult male *Kptn* mice (n=8 *Kptn*$^{-/-}$ versus 8 *Kptn*$^{+/+}$ controls). Below, a representative heatmap of the p-values for the three studied sections (Bregma +0.98mm, Bregma -1.34mm and Bregma -5.80mm). Above, histograms of the percentage change of *Kptn*$^{-/-}$ relative to *Kptn*$^{+/+}$ controls (100%). **b** Neuroanatomical findings in adult female *Kptn* mice (n=8 *Kptn*$^{-/-}$ versus 8 *Kptn*$^{+/+}$ controls), as described for males in **a**. In the legends in **c**, green text indicates length measurements and black text denotes area measurements. **d** Neuroanatomical findings in three week old (P20) male *Kptn* mice (n=6 *Kptn*$^{-/-}$ versus 6 *Kptn*$^{+/+}$ controls), as described for adults in **a**, with legend in **c**. **e** At Birth (P0), the total brain area (tba) of Kptn$^{-/-}$ mice (n=9) is unchanged (2.77% increase, p=0.494) compared to *Kptn*$^{+/+}$ controls (n=8). Only the hippocampus (11% increase, p=0.0489) and internal capsule (8.6% decrease, p=0.0361) show statistically significant changes at this stage of development (See Supplementary Table 2 for details). All p-values are from 2-sample two-tailed Student's t-tests.

**SUPPLEMENTARY FIGURE 5 Progressive macrocephaly in a representative KRD proband.** Occipitofrontal circumference in centimeters for a male proband (Proband 14, Supplementary Table 3) with KRD. OFC increased from below the 50$^{th}$ centile (mean) at birth to over two standard deviations (SD) above the mean by the age of two years (blue line). Centiles given in brackets.

**SUPPLEMENTARY FIGURE 6 Connectivity Map search identifies mTOR inhibitors as the most likely effective treatment for KRD.** Top negatively-ranked compounds identified by querying the Connectivity Map database with a differentially expressed gene set from *KPTN*$^{-/-}$ NPCs. **a** Torin-2 is identified as the strongest inversely-correlated compound across all cell types. **b** Query results of the Connectivity Map specifically in HCC515 cells identifies five known mTOR inhibitors among the top 32 negatively ranked compounds.

**SUPPLEMENTARY FIGURE 7 KRD models show dysregulation of numerous seizure associated genes. a** Mouse brain and **b** human NPC models of KRD show significant dysregulation of genes associated with disorders involving seizures (statistically significant $Log_2$-fold changes in expression are indicated by non-grey adjusted p-value heatmap cells). **c** Statistically significant overlap between mouse and human NPC seizure gene signatures (107 genes in common, 1.5 fold over-enriched, $p=3.6x10^{-7}$, hypergeometric test) in KRD models.

### SUPPLEMENTARY TABLE LEGENDS

**SUPPLEMENTARY TABLE 1 Microcomputed tomography findings in *Kptn*$^{+/+}$ and *Kptn*$^{-/-}$ mice reveal significant change to brain cavity dimensions in KRD model.** Inter-landmark distances were calculated (landmarks in Supplementary Figure 3, see Methods) and compared to identify statistically significant differences in mean parameter lengths between male **a** *Kptn*$^{+/+}$ and **b** *Kptn*$^{-/-}$ mice (n=5 each). Statistically significant p-values ($p<0.05$, 2-sample two-tailed Student's t-test) are indicated in green in **d.** All significant changes (boxed in c) show an increase in brain cavity size in *Kptn*$^{-/-}$ mice.

**SUPPLEMENTARY TABLE 2 Quantification of neuroanatomical features in the KRD mouse model at P0, P20, and adult stages.** Measurements of length or area of 53 features at birth and 78 features in P20 and adult brains were made (See Methods). **TAB 2A** is a list of the 78 neuroanatomical features for brain coronal analysis in P20 and adult stages. **TAB 2B** is the raw measurements for adult male mice at 16 weeks of age across all 78 parameters. **TAB 2C** is the raw measurements for adult female mice at 16 weeks of age across all 78 parameters. **TAB 2D** is the raw measurements for adult male mice at 16 weeks of cellular parameters including cell count, total cell area, cell circularity and cell solidity. **TAB 2E** is a list of the 53 neuroanatomical features for brain coronal analysis in P0 mice. **TAB 2F** is the raw measurements for mice at P0 across all 53 parameters. **TAB 2G** is the raw measurements for male mice mice at P20 across all 78 parameters. All p-values are from 2-sample two-tailed Student's t-tests.

**SUPPLEMENTARY TABLE 3 Occipitofrontal circumference measurements of KRD probands and parents.** Blue boxes: OFC measurements in centimeters (z-score in brackets) and genotype for previously published and newly identified individuals with KRD from birth to last reported measurement. Green boxes: OFC measurements in centimeters (z-score in brackets) and genotype for heterozygous parents of individuals with KRD. N/K = not known, WT = wild-type. *Progressive chart ID* indicates individuals plotted in Fig. 5b. *Family ID* identifies groups of siblings within nuclear families. *Reference* indicates first publication of individual. See Methods for details of z-score calculations.

**SUPPLEMENTARY TABLE 4 RNA-Seq analysis of differential gene expression in KRD models. TAB a**: DESEQ2 summary data across all mouse tissues and stages. **TAB b**: DESEQ2 summary data from *KPTN*$^{+/-}$ *and KPTN*$^{-/-}$ NPCs. *BaseMean* is normalised mean read count across the tissue for each gene, *log2FoldChange* is the Log2Fold change when comparing LoF model to matched wildtype samples, *padj* is the adjusted p-value output from DESEQ2 for the differential expression (see methods for details), *E18* denotes embryonic day 18 brain, *FrC* is prefrontal cortex, *Hipp* is hippocampus, *Cer* is cerebellum, *p21* denotes postnatal day 21, and *adult* denotes adult mice.

**SUPPLEMENTARY TABLE 5 KRD mouse model shows increased expression of progenitor markers.** Differential gene expression analysis between *KPTN*$^{+/+}$ and *KPTN*$^{-/-}$ mice reveals an increase in markers for neural stem cells (Radial Glia-like cells) and neurogenic intermediate progenitors (Intermediate Progenitors) at both P21 and in adults. Values are expressed as change compared to *KPTN*$^{+/+}$wild-type expression levels. All reported changes have adjusted p-values <0.05.

**SUPPLEMENTARY TABLE 6 Loss of KPTN function in human neural stem cells results in dysregulation of developmentally important haploinsufficient disorder-associated genes.** Twelve chromatin modifying transcriptional regulators, whose heterozygous loss-of-function in humans causes severe developmental disorders, are downregulated by 25-46% in *KPTN*$^{-/-}$ NPCs (*KPTN*$^{-/-}$ expression shown as % of *KPTN*$^{+/+}$ level, adjusted p-values<0.05). Phenotypes associated with loss of function in these disorders are relevant to those seen in KRD patients, including intellectual disability and/or developmental delay (ID/DD), seizures, autistic features, language deficits, and craniofacial dysmorphisms. Their presence was scored from published reports (References) as Present frequently (Y), occasionally (+/-), rarely (rare) or not observed (N).

**SUPPLEMENTARY FILE**

**SUPPLEMENTARY FILE 1 Gene enrichment in mouse KRD model RNA-Seq. Tab a,** Summary table of mTOR pathway relevant Gene Ontology enrichment findings for mouse RNA-Seq samples. **Tab b,** Results of hypergeometric test specifically for enrichment of mTOR pathway genes in the KRD model mouse RNA-Seq samples ($\alpha$<0.05). Complete individual Gene Ontology enrichment findings are provided in labelled tabs.
