## Supplementary Fig. 1 for "Mouse and cellular models of *KPTN*-related disorder implicate mTOR signalling in cognitive and progressive overgrowth phenotypes"

Supplementary Figure 1

**a**

```
Exon 8(NM_007059.4): AGGTTCTGCAGATGTGGTCGGTCCTGCAGGACGGTCCCATCTCCCGAGTGATTGTGTTCAGCCTCTCGGCCGCCAAGG
c.714_731dup:        AGGTTCTGCAGATGTGGTCGGTCCTGCAGGACGGTCCCATCTCCCGAGTGATTGTGTTCAGCCTCTCGGCCGCCAAGG
c.776C>A:            AGGTTCTGCAGATGTGGTCGGTCCTGCAGGACGGTCCCATCTCCCGAGTGATTGTGTTCAGCCTCTAGGCCGCCAAGG
KPTN -/- allele 1:   AGGT-------------------------GACGGTCCC-------GAGTGATTGTGTTCAGCCTCTCGGCCGCCAAGG
KPTN -/- allele 2:   AGGTTCTGCAGATGTGGTCGGTCCTGCAGGACGGTCCC-------GAGTGATTGTGTTCAGCCTCTCGGCCGCCAAGG
KPTN +/- allele 1:   AGGTTCTGCAGATGTGGTCGGTCC--------------------CGAGTGATTGTGTTCAGCCTCTCGGCCGCCAAGG
```

**b**

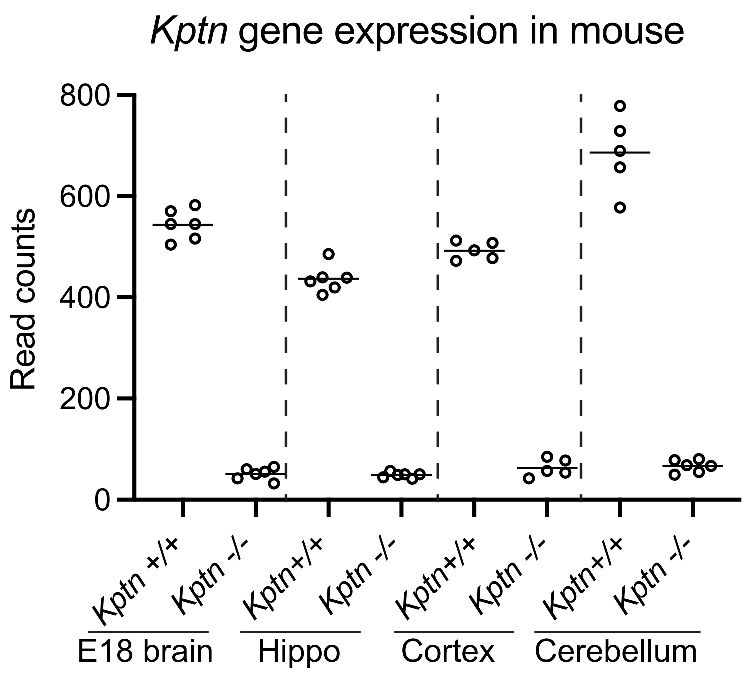


**c**

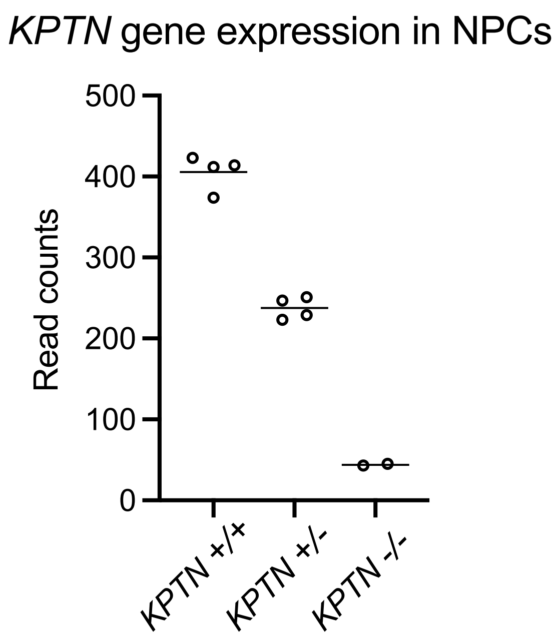
