## Supplementary figures and images for "Mouse and cellular models of *KPTN*-related disorder implicate mTOR signalling in cognitive and progressive overgrowth phenotypes"

### Supplementary Fig. 2

Supplementary Figure 2

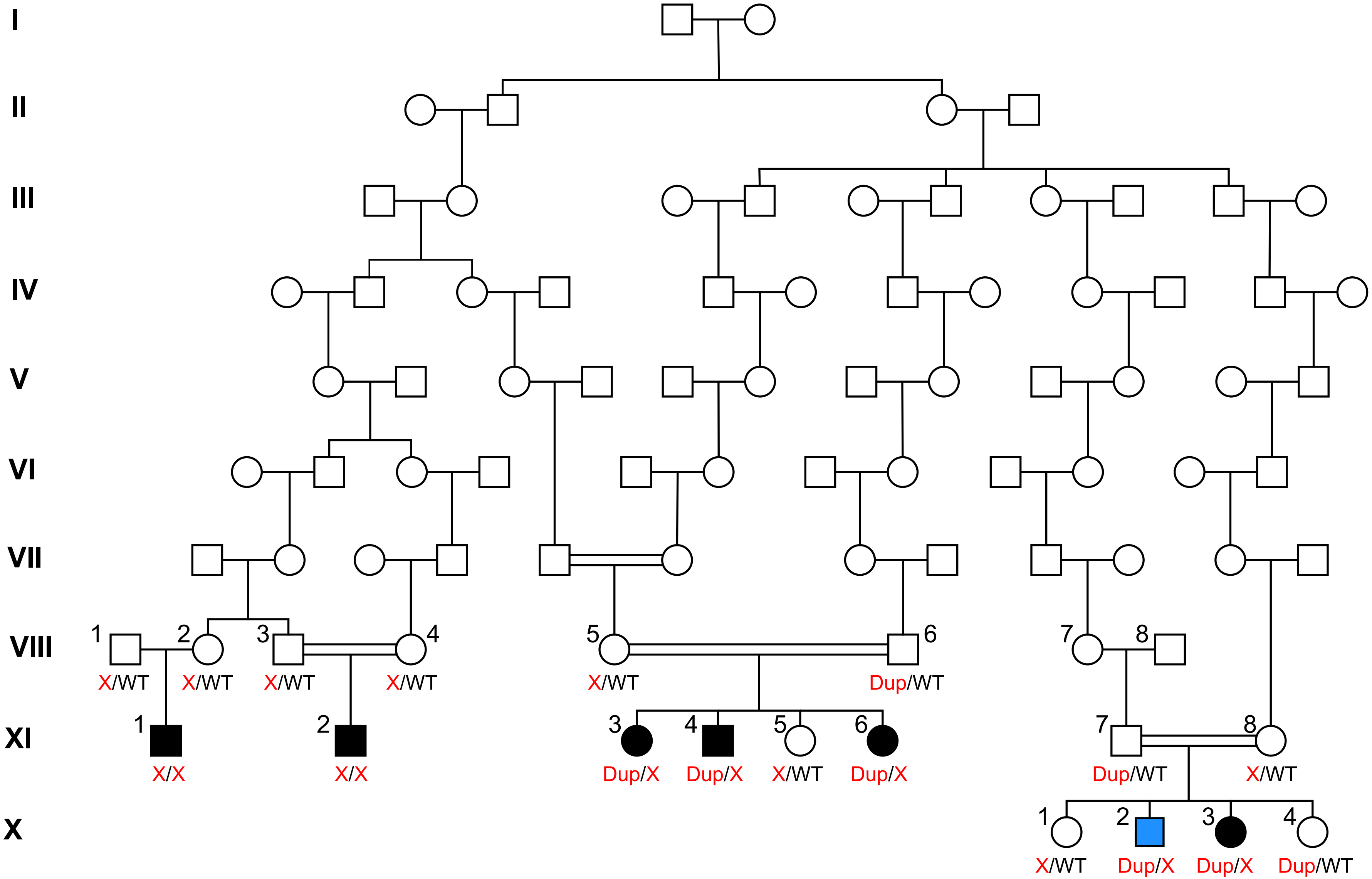

### Supplementary Fig. 5

Supplementary Figure 5

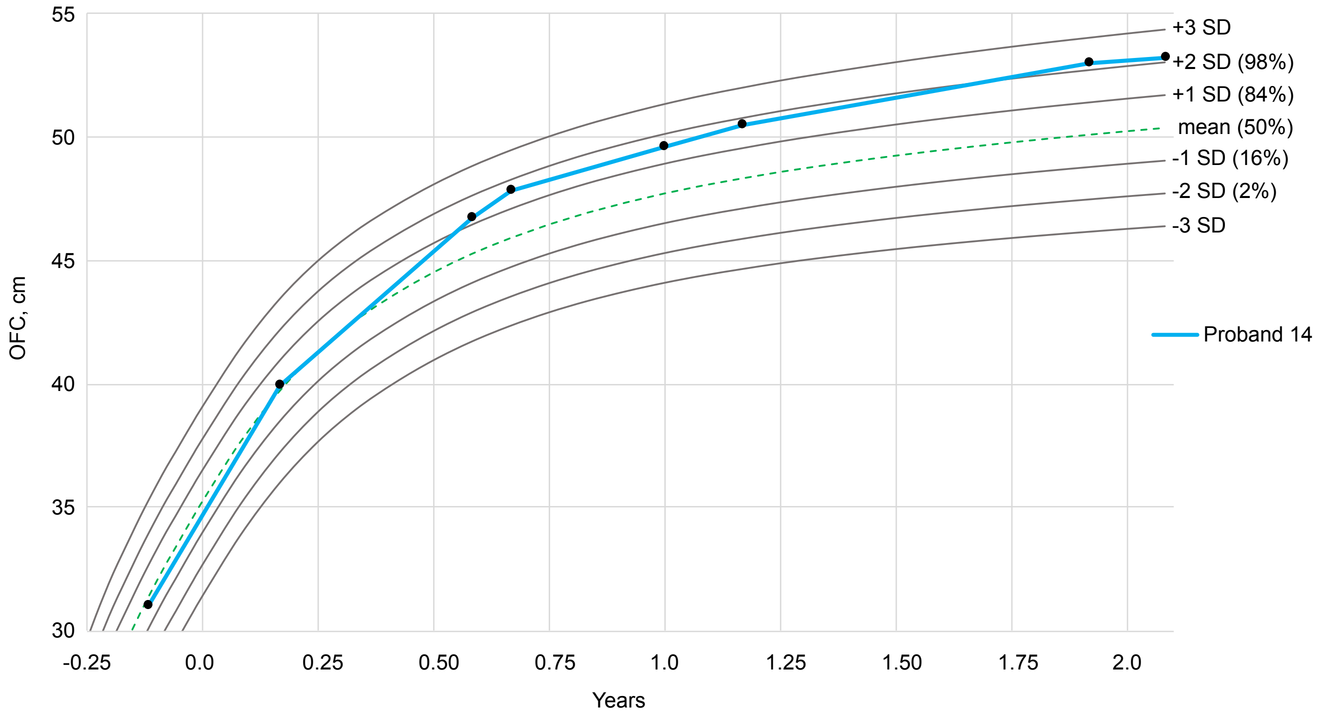

### Supplementary Fig. 7

#### Supplementary Figure 7

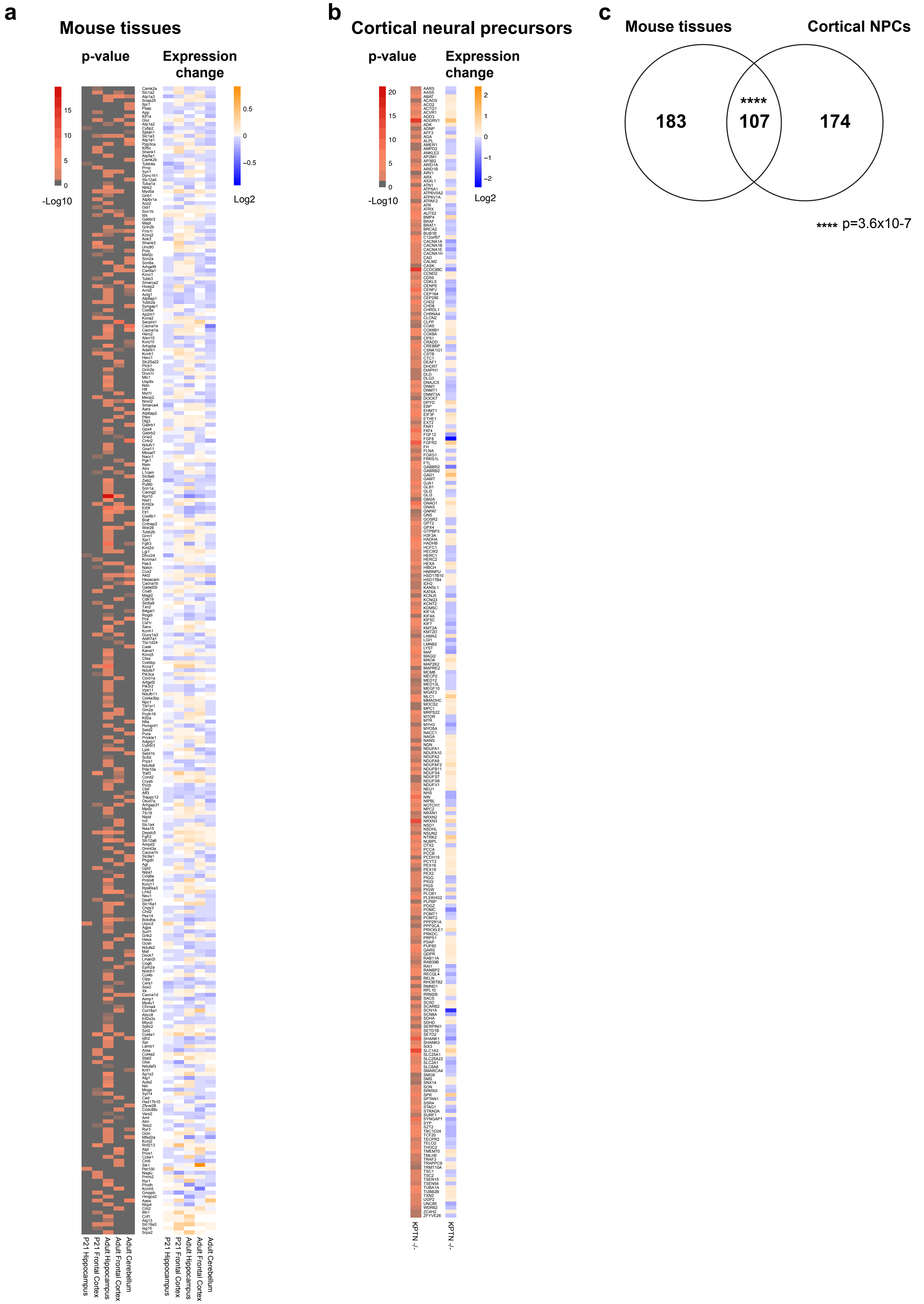
