## Supplementary Fig. 3 for "Mouse and cellular models of *KPTN*-related disorder implicate mTOR signalling in cognitive and progressive overgrowth phenotypes"

Supplementary Figure 3

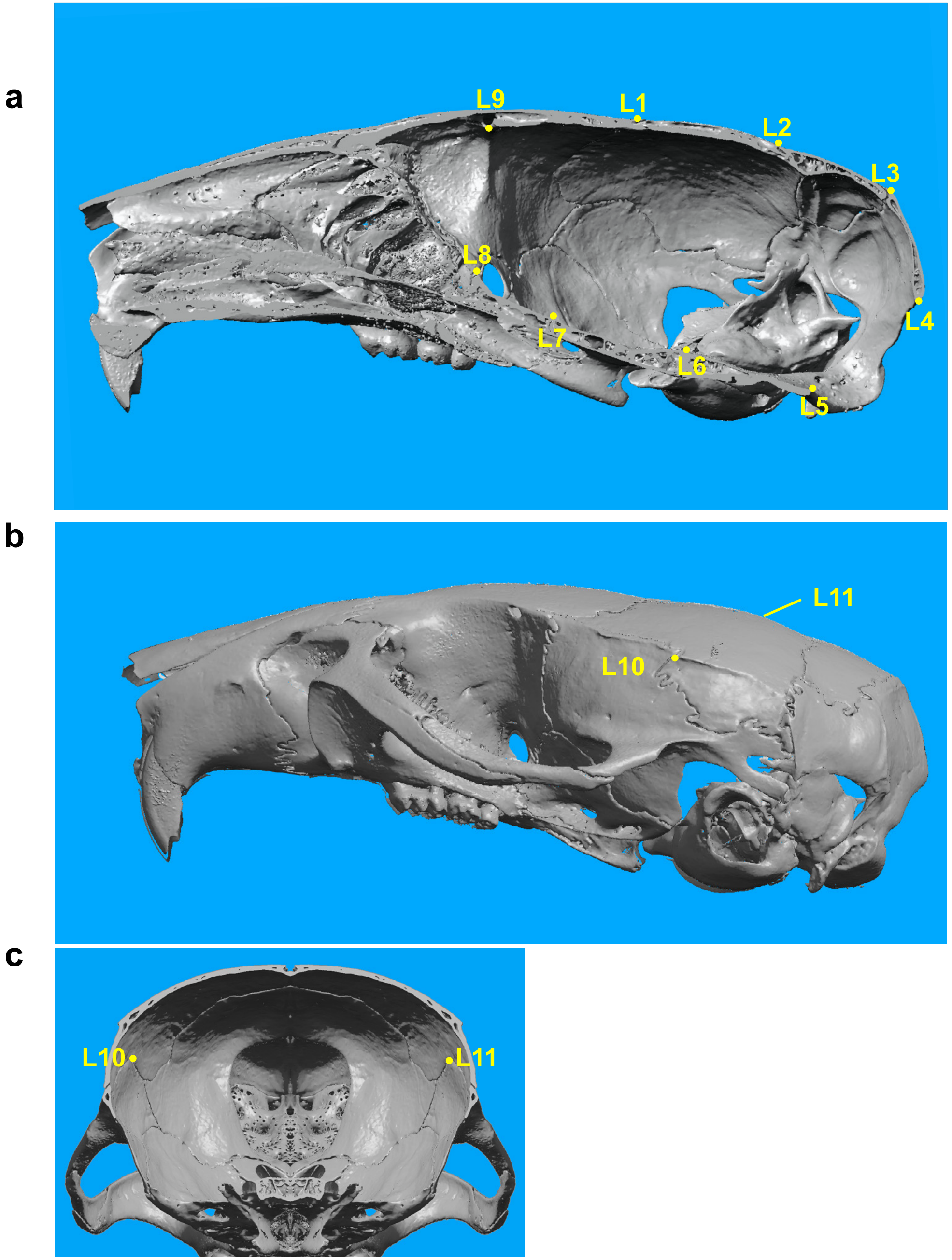


| | |
|---|---|
| L1 | Bregma |
| L2 | Lambda |
| L3 | Intersection of interparietal bones with squamous portion of occipital bone at midline |
| L4 | Opisthion, midsagittal point on the posterior margin of the foramen magnum |
| L5 | Caudal most point of basi-occipital bone at mid-sagittal plane |
| L6 | Dorsal-most point of spheno-occipital synchondrosis at mid-sagittal plane |
| L7 | Dorsal-most point of inter-sphenoid synchondrosis at mid-sagittal plane |
| L8 | Rostral end of pre-sphenoid bone at mid-sagittal plane |
| L9 | Caudal-most point of the roof of the olfactory fossa |
| L10 | Intersection of the saquamosal suture with temporal crest, Left |
| L11 | Intersection of the saquamosal suture with temporal crest, Right |
