## Supplementary Fig. 4 for "Mouse and cellular models of *KPTN*-related disorder implicate mTOR signalling in cognitive and progressive overgrowth phenotypes"

Supplementary Figure 4

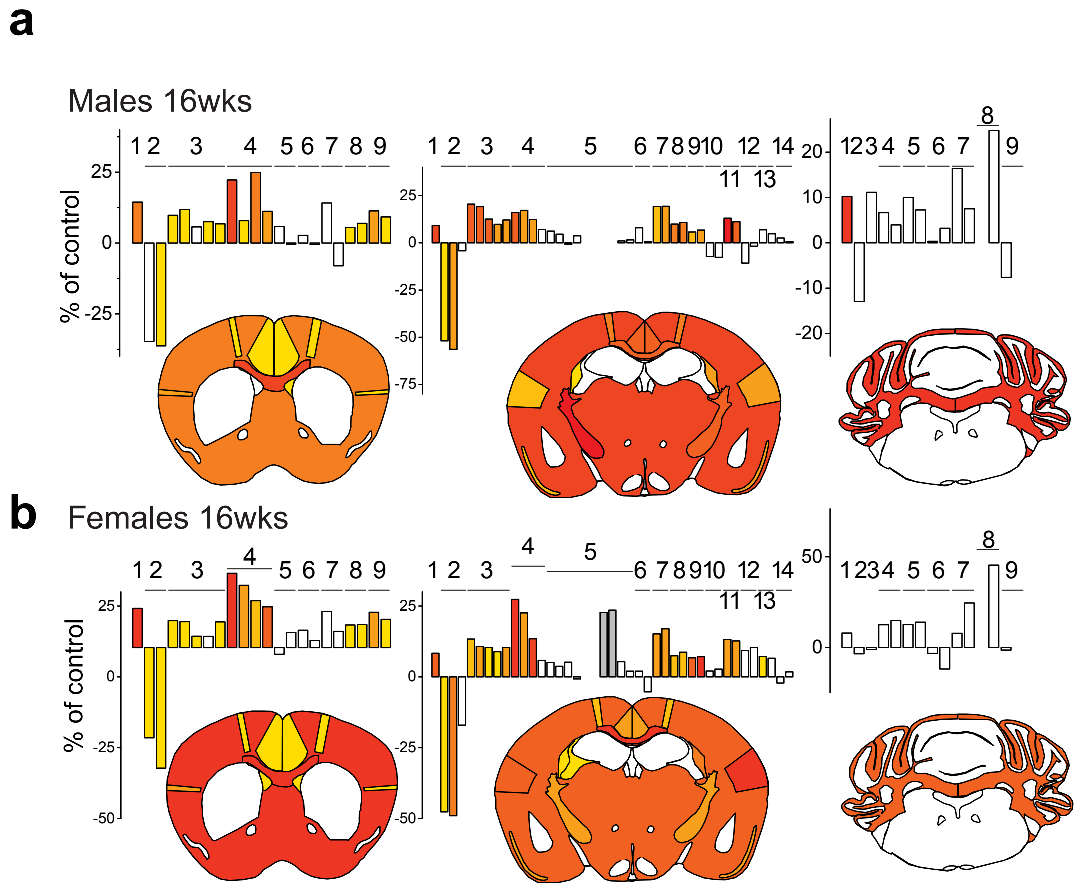


c

**Section1**

1 Total Brain Area
2 Lateral Ventricle_Left side
Lateral Ventricle_Right side
3 Cingulate cortex area_Left side
Cingulate cortex area_Right side
Cingulate cortex area_Width_Left side
Cingulate cortex_Width_Right side
Cingulate cortex_Height
4 genu of the corpus callosum
genu of the corpus callosum_Height
genu of the corpus callosum_Width_Basal
genu of the corpus callosum_Width_Top
5 caudate putamen_Left side
caudate putamen_Right side
6 anterior_commissure_Left side
anterior_commissure_Right side
7 piriform cortex_Left side
piriform cortex_Right side
8 primary motor cortex_Left side length
primary motor cortex_Reft side lengthcc
9 secondary somatosensory cortex_Left side length
secondary somatosensory cortex_Right side length

**Section 3**

1 Total Brain Area
2 fourth Ventricle
3 pons
4 pyramidal_L
pyramidal_R
5 genu facial nerve_L
genu facial nerve_R
6 cochlear nucleus_L
cochlear nucleus_R
7 lateral dentate Cb_L
lateral dentate Cb_R
8 interposed cerebellar_L
interposed cerebellar_R
9 internal granule layer
folia number

**Section 2**

1 Total Brain Area
2 Lateral Ventricle_Left side
Lateral Ventricle_Right side
dorsal 3rd ventricle
3 retrosplenial granular cortex_Left side
retrosplenial granular cortex_Right side
retrosplenial granular cortex_Width_Left side
retrosplenial granular cortex_Width_Right side
retrosplenial granular cortex_Height
4 corpus callosum
corpus callosum_Height
ccorpus callosum_Width
dorsal hippocampal commissure
5 hippocampus
pyramidal layer
dentate gyrus_Left side
dentate gyrus_Right side
lacunosum moleculare_Left side
lacunosum moleculare_Right side
radiatum layer_Left side
radiatum layer_Right side
oriens layer_Left side
oriens layer_R
6 amygdala_Left side
amygdala_Right side
7 piriform cortex_Left side
piriform cortex_Right side
8 primary motor cortex_Left side length
primary motor cortex_Right side length
9 secondary somatosensory cortex_Left side length
secondary somatosensory cortex_Right side length
10 mammillothalamic tract_Left side
mammillothalamic tract_Right side
11 internal capsule_Left side
internal capsule_Right side
12 optic tract_Left side
optic tract_Right side
13 fimbria_Left side
fimbria_Right side
14 med habenular_Left side
med habenular_Right side

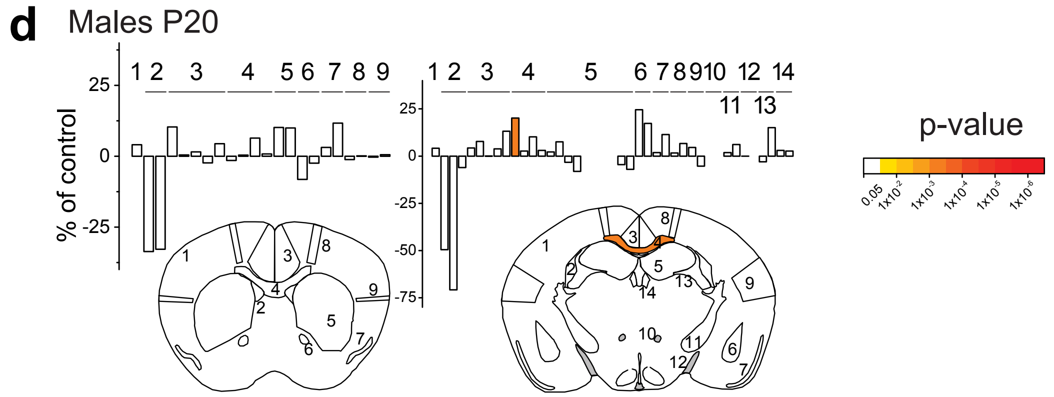


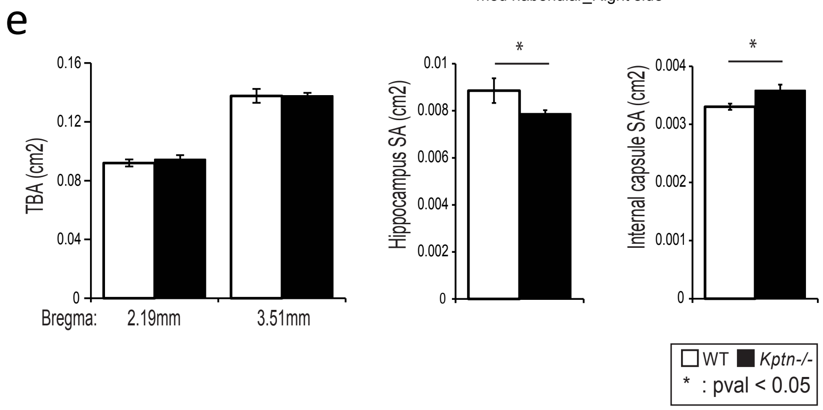
