## Supplementary Fig. 6 for "Mouse and cellular models of *KPTN*-related disorder implicate mTOR signalling in cognitive and progressive overgrowth phenotypes"

Supplementary figure 6

**a**

Connectivity Map overall scores across all cell lines

| Rank | Score | ID | Compound | Description | Target |
|---|---|---|---|---|---|
| 1 | -96.28 | BRD-K68174511 | torin-2 | mTOR inhibitor | MTOR |
| 2 | -91.63 | BRD-A99449986 | MT-21 | Caspase activator | CYCS, SLC25A4 |
| 3 | -84.80 | BRD-K37798499 | etoposide | Topoisomerase inhibitor | TOP2A, CYP2E1, CYP3A5, TOP2B |
| 4 | -82.27 | BRD-A25687296 | emetine | Protein synthesis inhibitor | RPS2 |
| 5 | -81.30 | BRD-K36395411 | SB-206553 | Serotonic receptor antagonist | HTR2B, HTR2C, HTR1A, HTR2A |
| 6 | -80.70 | BRD-K02130563 | panobinostat | HDAC inhibitor | HDAC1, HDAC2, HDAC3, HDAC4, HDAC6, HDAC7, HDAC8, HDAC9 |

**b**

HCC515 cell line

| Score | Rank | Compound | Description |
|---|---|---|---|
| -99.56 | 1 | torin-2 | mTOR inhibitor |
| -99.04 | 3 | QL-X-138 | mTOR inhibitor |
| -95.89 | 12 | torin-1 | mTOR inhibitor |
| -93.44 | 30 | sirolimus | mTOR inhibitor |
| -93.23 | 32 | WYE-354 | mTOR inhibitor |
