## Supplementary Table 1 for "Mouse and cellular models of *KPTN*-related disorder implicate mTOR signalling in cognitive and progressive overgrowth phenotypes"

**a** Mean distance, Wild type

| | L2 | L3 | L4 | L5 | L6 | L7 | L8 | L9 | L11 | | |
|---|---|---|---|---|---|---|---|---|---|---|---|
| L1 | 4.275 | 7.495 | 9.344 | 8.967 | 6.560 | 5.770 | 5.999 | 3.456 | | L1 | Bregma |
| L2 | | 3.283 | 5.587 | 6.674 | 6.187 | 7.728 | 9.085 | 7.682 | | L2 | Lambda |
| L3 | | | 2.890 | 5.662 | 7.038 | 9.727 | 11.555 | 10.823 | | L3 | Intersection of interparietal bones with squamous portion of occipital bone at midline |
| L4 | | | | 3.781 | 6.541 | 9.925 | 12.119 | 12.361 | | L4 | Opisthion, midsagittal point on the posterior margin of the foramen magnum |
| L5 | | | | | 3.662 | 7.318 | 9.738 | 11.201 | | L5 | Caudal most point of basi-occipital bone at mid-sagittal plane |
| L6 | | | | | | 3.659 | 6.083 | 8.005 | | L6 | Dorsal-most point of spheno-occipital synchondrosis at mid-sagittal plane |
| L7 | | | | | | | 2.455 | 5.496 | | L7 | Dorsal-most point of inter-sphenoid synchondrosis at mid-sagittal plane |
| L8 | | | | | | | | 4.248 | | L8 | Rostral end of pre-sphenoid bone at mid-sagittal plane |
| L9 | | | | | | | | | | L9 | Caudal-most point of the roof of the olfactory fossa |
| L10 | | | | | | | | | 9.5528 | L10 | Intersection of the saquamosal suture with temporal crest |

**b** Mean distance, *Kptn* -/-

| | L2 | L3 | L4 | L5 | L6 | L7 | L8 | L9 | L11 | | |
|---|---|---|---|---|---|---|---|---|---|---|---|
| L1 | 4.235 | 7.467 | 9.581 | 9.183 | 6.673 | 6.072 | 6.262 | 3.721 | | L1 | Bregma |
| L2 | | 3.310 | 5.938 | 6.986 | 6.264 | 7.904 | 9.220 | 7.885 | | L2 | Lambda |
| L3 | | | 3.227 | 5.898 | 7.000 | 9.779 | 11.599 | 11.005 | | L3 | Intersection of interparietal bones with squamous portion of occipital bone at midline |
| L4 | | | | 3.807 | 6.483 | 9.963 | 12.197 | 12.707 | | L4 | Opisthion, midsagittal point on the posterior margin of the foramen magnum |
| L5 | | | | | 3.622 | 7.256 | 9.707 | 11.428 | | L5 | Caudal most point of basi-occipital bone at mid-sagittal plane |
| L6 | | | | | | 3.653 | 6.088 | 8.142 | | L6 | Dorsal-most point of spheno-occipital synchondrosis at mid-sagittal plane |
| L7 | | | | | | | 2.485 | 5.706 | | L7 | Dorsal-most point of inter-sphenoid synchondrosis at mid-sagittal plane |
| L8 | | | | | | | | 4.365 | | L8 | Rostral end of pre-sphenoid bone at mid-sagittal plane |
| L9 | | | | | | | | | | L9 | Caudal-most point of the roof of the olfactory fossa |
| L10 | | | | | | | | | 9.797 | L10 | Intersection of the saquamosal suture with temporal crest |

**c** Difference in length, as % of wildtype

| | L2 | L3 | L4 | L5 | L6 | L7 | L8 | L9 | L11 | | |
|---|---|---|---|---|---|---|---|---|---|---|---|
| L1 | -0.9 | -0.4 | 2.5 | 2.4 | 1.7 | 5.2 | 4.4 | 7.7 | | L1 | Bregma |
| L2 | | 0.8 | 6.3 | 4.7 | 1.2 | 2.3 | 1.5 | 2.6 | | L2 | Lambda |
| L3 | | | 11.7 | 4.2 | -0.5 | 0.5 | 0.4 | 1.7 | | L3 | Intersection of interparietal bones with squamous portion of occipital bone at midline |
| L4 | | | | 0.7 | -0.9 | 0.4 | 0.6 | 2.8 | | L4 | Opisthion, midsagittal point on the posterior margin of the foramen magnum |
| L5 | | | | | -1.1 | -0.9 | -0.3 | 2.0 | | L5 | Caudal most point of basi-occipital bone at mid-sagittal plane |
| L6 | | | | | | -0.2 | 0.1 | 1.7 | | L6 | Dorsal-most point of spheno-occipital synchondrosis at mid-sagittal plane |
| L7 | | | | | | | 1.2 | 3.8 | | L7 | Dorsal-most point of inter-sphenoid synchondrosis at mid-sagittal plane |
| L8 | | | | | | | | 2.8 | | L8 | Rostral end of pre-sphenoid bone at mid-sagittal plane |
| L9 | | | | | | | | | | L9 | Caudal-most point of the roof of the olfactory fossa |
| L10 | | | | | | | | | 2.6 | L10 | Intersection of the saquamosal suture with temporal crest |

**d** t-test p-values

| | L2 | L3 | L4 | L5 | L6 | L7 | L8 | L9 | L11 | | |
|---|---|---|---|---|---|---|---|---|---|---|---|
| L1 | 0.6483 | 0.8313 | 0.0684 | 0.034 | 0.2241 | 0.0017 | 0.003 | 0.0042 | | L1 | Bregma |
| L2 | | 0.7947 | 0.0285 | 0.0055 | 0.438 | 0.0686 | 0.1884 | 0.0196 | | L2 | Lambda |
| L3 | | | 0.0822 | 0.0199 | 0.5744 | 0.4949 | 0.6458 | 0.0825 | | L3 | Intersection of interparietal bones with squamous portion of occipital bone at midline |
| L4 | | | | 0.7032 | 0.2547 | 0.5696 | 0.3133 | 0.0125 | | L4 | Opisthion, midsagittal point on the posterior margin of the foramen magnum |
| L5 | | | | | 0.4105 | 0.2673 | 0.5179 | 0.0278 | | L5 | Caudal most point of basi-occipital bone at mid-sagittal plane |
| L6 | | | | | | 0.8701 | 0.9027 | 0.0428 | | L6 | Dorsal-most point of spheno-occipital synchondrosis at mid-sagittal plane |
| L7 | | | | | | | 0.3858 | 0.0084 | | L7 | Dorsal-most point of inter-sphenoid synchondrosis at mid-sagittal plane |
| L8 | | | | | | | | 0.0849 | | L8 | Rostral end of pre-sphenoid bone at mid-sagittal plane |
| L9 | | | | | | | | | | L9 | Caudal-most point of the roof of the olfactory fossa |
| L10 | | | | | | | | | 0.0356 | L10 | Intersection of the saquamosal suture with temporal crest |
