## Supplementary Table 5 for "Mouse and cellular models of *KPTN*-related disorder implicate mTOR signalling in cognitive and progressive overgrowth phenotypes"

| Ensembl ID | Gene | P21 Frontal cortex | Hippocampus | Frontal cortex | Cerebellum | Cell types |
|---|---|---|---|---|---|---|
| ENSMUSG00000004891 | Nes | | | 13.7% | | Radial Glia-like cells |
| ENSMUSG00000026728 | Vim | | | 11.6% | | |
| ENSMUSG00000005360 | GLAST/Slc1a3 | 8.7% | -8% | 7.8% | | |
| ENSMUSG00000035686 | Thrsp | 51.1% | 13.5% | 38.8% | | |
| ENSMUSG00000027004 | Frzb | | 25.2% | | | |
| ENSMUSG00000059325 | Hopx | | | 17.8% | | |
| ENSMUSG00000032446 | Eomes | | | | 61% | Intermediate Progenitors |
| ENSMUSG00000035033 | Tbr1 | | 10% | | | |
| ENSMUSG00000038255 | Neurod2 | | 16.5% | | | |
| ENSMUSG00000020423 | Btg2 | | | 49.3% | | |
| ENSMUSG00000076431 | Sox4 | | 8.9% | | | |
