## Supplementary Table 6 for "Mouse and cellular models of *KPTN*-related disorder implicate mTOR signalling in cognitive and progressive overgrowth phenotypes"

Phenotypes of haploinsufficient LoF genes dysregulated in *KPTN*-/- NPCs

| Gene | ID/DD | Haplo-insufficient LOF | Expression in KPTN-/- NPCs, % of wildtype | Craniofacial phenotypes | Head size | Seizures | Autistic features | Language/speech problems | Brain/nervous system structural abnormalities | References |
|---|---|---|---|---|---|---|---|---|---|---|
| AUTS2 | Y | Y | 71.3% | Y | Y (smaller) | Y | Y | Y | +/- | PMID: 23332918, PMID: 27075013 |
| AHDC1 | Y | Y | 72.4% | Y | +/- (some macro) | +/- | Y | Y | Y | PMID: 24791903, PMID: 27148574 |
| ANKRD11 | Y | Y | 56.8% | Y | rare (smaller) | +/- | +/- | +/- | Y | PMID: 21782149, PMID: 31191201 |
| CHD7 | Y | Y | 72.9% | Y | N | N | +/- | (hearing loss) | Y | PMID: 16400610, PMID: 15300250, PMID: 16155193 |
| CREBBP | Y | Y | 73.5% | Y | Y (smaller) | +/- | +/- | Y | N | PMID: 12070251, PMID: 18792986, PMID: 26788536 |
| CHD2 | Y | Y | 73.5% | N | N | Y | +/- | N | +/- | PMID: 23708187, PMID: 25672921 |
| EBF3 | Y | Y | 53.7% | Y | N | rare | N | Y | N | PMID: 29062322, PMID: 28017373 |
| KMT2D | Y | Y | 75.8% | Y | Y (smaller) | Y | +/- | Y | Y | PMID: 21671394, PMID: 21607748, PMID: 15578615 |
| TBR1 | Y | Y | 71.3% | N | N | N | Y | Y | Y | PMID: 25232744, PMID: 30268909, PMID: 29288087 |
| TCF20 | Y | Y | 75.2% | Y | Y (some macro) | +/- | Y | Y | +/- | PMID: 30819258 |
| SETD1B | Y | Y | 74.9% | Y | N | Y | Y | Y | +/- | PMID: 29322246, PMID: 31110234, PMID: 31685013, PMID: 32546566 |
| ZIC2 | Y | Y | 71.5% | Y | N | N | N | N | Y | PMID: 9771712, PMID: 11285244, PMID: 19955556 |
